# The *C. elegans* Myc-family of transcription factors coordinate a dynamic adaptive response to dietary restriction

**DOI:** 10.1101/2023.11.22.568222

**Authors:** Adam Cornwell, Yun Zhang, Manjunatha Thondamal, David W. Johnson, Juilee Thakar, Andrew V. Samuelson

## Abstract

Dietary restriction (DR), the process of decreasing overall food consumption over an extended period of time, has been shown to increase longevity across evolutionarily diverse species and delay the onset of age-associated diseases in humans. In *Caenorhabditis elegans*, the Myc-family transcription factors (TFs) MXL-2 (Mlx) and MML-1 (MondoA/ChREBP), which function as obligate heterodimers, and PHA-4 (orthologous to forkhead box transcription factor A) are both necessary for the full physiological benefits of DR. However, the adaptive transcriptional response to DR and the role of MML-1::MXL-2 and PHA-4 remains elusive. We identified the transcriptional signature of *C. elegans* DR, using the *eat-2* genetic model, and demonstrate broad changes in metabolic gene expression in *eat-2* DR animals, which requires both *mxl-2* and *pha-4*. While the requirement for these factors in DR gene expression overlaps, we found many of the DR genes exhibit an opposing change in relative gene expression in *eat-2;mxl-2* animals compared to wild-type, which was not observed in *eat-2* animals with *pha-4* loss. We further show functional deficiencies of the *mxl-2* loss in DR outside of lifespan, as *eat-2;mxl-2* animals exhibit substantially smaller brood sizes and lay a proportion of dead eggs, indicating that MML-1::MXL-2 has a role in maintaining the balance between resource allocation to the soma and to reproduction under conditions of chronic food scarcity. While *eat-2* animals do not show a significantly different metabolic rate compared to wild-type, we also find that loss of *mxl-2* in DR does not affect the rate of oxygen consumption in young animals. The gene expression signature of *eat-2* mutant animals is consistent with optimization of energy utilization and resource allocation, rather than induction of canonical gene expression changes associated with acute metabolic stress -such as induction of autophagy after TORC1 inhibition. Consistently, *eat-2* animals are not substantially resistant to stress, providing further support to the idea that chronic DR may benefit healthspan and lifespan through efficient use of limited resources rather than broad upregulation of stress responses, and also indicates that MML-1::MXL-2 and PHA-4 may have different roles in promotion of benefits in response to different pro-longevity stimuli.

## Introduction

Dietary restriction (DR) is one of few known interventions that results in delayed onset of aging-associated disease and improves longevity in a broad range of evolutionarily distinct animals ranging from invertebrates to monkeys, and recently has been found to improve health biomarkers in humans (Kennedy et al. 2007; Fontana et al. 2010; Nakagawa et al. 2012; Kapahi et al. 2017; Mattison et al. 2017; Il’yasova et al. 2018; Kraus et al. 2019; Krittika and Yadav 2019; von Frieling and Roeder 2020; Souza Matos et al. 2021; Green et al. 2022). DR is a broad term encompassing a growing number of treatment paradigms, including chronic restriction of overall energy intake or reduction of specific nutrients (*e.g.* methionine, carbohydrates) and intermittent fasting: scheduled short-term food withdrawal as with alternate-day feeding or time-restricted feeding (Greer and Brunet 2009; Soultoukis and Partridge 2016; Duregon et al. 2021). Recent clinical trials in humans have found significant improvement in biomarkers of cardiovascular health in both short term alternate-day fasting (Stekovic et al. 2019) and reduced average energy intake reduction over several years (Il’yasova et al. 2018; Kraus et al. 2019). Although the benefits of DR on organismal health are widely conserved, the full mechanistic picture of how various types of DR influences physiological decline during aging remains far from clear, due to complex interactions between genetics, diet, and environment that influence aging(Mair and Dillin 2008).

*C. elegans* is a preeminent model for aging studies, in part due to its normally short lifespan, fast generation time, and wealth of genetic tools. Working with an intact metazoan model enables identification of cell non-autonomous interactions across distinct tissues, including those coordinated by a nervous system (Brenner and Brenner 1974). Adult wild-type hermaphrodite animals have 959 somatic cells that arise from an invariant and well-characterized lineage, without contribution of stem or progenitor cells to the maintenance of adult somatic tissues (Sulston and Horvitz 1977). This less-complex environment allows focus on elucidating genetic determinants of health and maintenance of differentiated somatic cells that act within an integrated biological system, without interference from regenerative cellular quality control mechanisms that preserve tissue homeostasis. Notably, the discovery of long-lived genetic mutants of *C. elegans* combined with the development of feeding-based RNAi has led to uncovering more than 1,000 genes that influence lifespan when perturbed, which we refer to as “gerogenes” (Johnson and Wood 1982; Johnson 1990; Kenyon et al. 1993; Timmons and Fire 1998; Murphy et al. 2003; Hamilton et al. 2005; Hansen et al. 2005; Samuelson et al. 2007; Lee et al. 2010; Bennett et al. 2014; Cornwell et al. 2018).

Many gerogenes are nodes in networks that sense nutrient status and are components of intricate networks connecting nutrient-associated signals to organismal longevity (Ogg et al. 1997; Jia et al. 2004; Hansen et al. 2005; Gelino et al. 2016; Gao et al. 2017; Denzel et al. 2018). For example, loss-of-function mutations in *daf-2*, orthologous to the insulin and IGF1 receptors, extends longevity through activation of the DAF-16 transcription factor (FOXO homolog) (Kenyon et al. 1993; Lin et al. 1997). Increasing DAF-16 expression is sufficient to extend longevity and improve survival to a myriad cellular stresses (Murphy et al. 2003; Mukhopadhyay et al. 2006). Studies into genetic variants associated with healthy aging in human populations have identified polymorphisms over-represented in human centenarians, such as FOXO3 a homolog of *daf-16* (Willcox et al. 2008). In addition to IIS, most gerogenes have evolutionarily conserved roles in the control of longevity including: the Target of Rapamycin (TOR) complexes, which regulate the balance between anabolism and catabolism based on amino acid and carbohydrate availability (Jia et al. 2004; Soukas et al. 2009; Lapierre et al. 2015; Edwards et al. 2015); and AMP kinase (AMPK), which responds to insufficient ATP levels (Apfeld et al. 2005; Greer et al. 2007; Greer and Brunet 2009; Weir et al. 2017). Overall, nearly 75% of gerogenes in *C. elegans* show considerable homology with mammals and humans, to the extent of functional substitution in some cases (Yuan et al. 1998; Harris et al. 2009; Cornwell et al. 2018; Kim et al. 2018; Tacutu et al. 2018). However, the majority of gerogenes remain under-studied, and have not been placed in an interaction network or pathway (Cornwell et al. 2018).

The *eat-2* genetic mutant is a common DR model in *C. elegans*; *eat-2* encodes a nicotinic acetylcholine receptor expressed in neuromuscular junctions of two pharyngeal muscle cells that contributes to control of pharyngeal pumping rate, thereby influencing food consumption (Mckay et al. 2004). Reduction of *eat-2* function results in decreased feeding rates and increased longevity, which provides a genetic model for studying the molecular and cellular basis of DR (Lakowski and Hekimi 1998; Mckay et al. 2004; Rodríguez-Palero et al. 2018). A number of transcriptional effectors of DR have been identified using this model, most notably the FOXA homolog PHA-4 (Panowski et al. 2007), which is specific as the FOXO homolog DAF-16 is not required (Lakowski and Hekimi 1998).

The MYC network of basic helix-loop-helix leucine zipper (bHLH Zip) transcription factors is perhaps best known for its namesake: MYC, which is a key regulator of metabolism, cell growth, and survival. MYC is also a well-characterized proto-oncogene that contributes to tumorigenic transformation in a range of cancers (Sloan and Ayer 2010; Conacci-Sorrell et al. 2014). MYC functionally regulates transcription when part of a heterodimeric complex with MAX. The requirement to bind MAX serves to regulate MYC activity, as MAX is competitively bound by MXD and MNT factors, which, in turn also compete for binding to MLX. These interactions further serve to regulate the activity of an additional set of metabolism-regulating transcription factors, MLXIP and MLXIPL (formerly known as MONDOA and MONDOB/CHREBP, respectively) (Conacci-Sorrell et al. 2014). All of these proteins form heterodimers at enhancer-box (E-box) sites (Sloan and Ayer 2010; Allevato et al. 2017; de Martin et al. 2021). The general properties of the MYC network factors are broadly conserved through metazoan evolution, although not all animals have a representative homolog for each of the core protein families in the network that are found in vertebrates (McFerrin and Atchley 2011). *C. elegans* in particular does not have a canonical MYC or MNT homolog, but does possess homologs for the other network members (McFerrin and Atchley 2011).

The MML-1::MXL-2 complex (mammalian MLXIP::MLX) in particular, and generally the entire *C. elegans* MYC network, increasingly appears to be a convergence point in multiple interaction networks involved in longevity and homeostatic maintenance. All of the Myc-family of transcription factors have been implicated in *C. elegans* longevity (*mdl-1, mxl-1, mml-1, mxl-2, mxl-3*) (O’Rourke and Ruvkun 2013; Johnson et al. 2014; Nakamura et al. 2016). MDL-1 (MXD) heterodimerizes with MXL-1 (MAX), loss of either *mdl-1* or *mxl-1* increases longevity and is dependent upon the MML-1::MXL-2 complex (Pickett et al. 2007; Johnson et al. 2014). Interestingly, in humans loss of a homolog of MML-1: MLXIPL, has been implicated in Williams-Beuren syndrome, a disease with progeric symptoms (Cairo et al. 2001) and reduced MYC expression increases longevity in mice (Hofmann et al. 2015). We previously demonstrated that the *C. elegans* MML-1::MXL-2 complex was necessary for the extended longevity conferred by chronic DR (*eat-2*) or reduced IIS; conversely loss of *mdl-1* or *mxl-1* does not further increase the longevity of these mutants (Samuelson et al. 2007; Johnson et al. 2014). *mml-1* and *mxl-2* are required for the extended longevity conferred via cell non-autonomous signals from the germline (*glp-1* mutant animals), after inactivation of TORC1, and increased activity of homeodomain-interacting protein kinase *hpk-1*, a neuronal transcriptional regulator of autophagy and longevity negatively regulated by TORC1 activity (Nakamura et al. 2016; Das et al. 2017; Lazaro-Pena et al. 2023). Given our previous finding that MML-1::MXL-2 is required for the longevity of *eat-2(ad465)* animals (Johnson et al. 2014), the importance of this complex in the integration of multiple pro-longevity signals, and the lack of known targets in the context of DR, we sought to define the adaptive transcriptional response to DR and the role of MML-1::MXL-2 therein (Johnson et al. 2014; Nakamura et al. 2016).

## Materials and Methods

### C. elegans strains

N2 Bristol was used as the wild-type strain, and all other strains were back-crossed at least six times to N2 before proceeding. The *mxl-2(tm1516)* strain was created by and obtained from Dr. Shohei Mitani with the Japanese Bioresource Project. Strains were confirmed to be WT for *fln-2(OT611)*, a background longevity mutation found in some commonly used wild-type strains (Zhao et al. 2019).

The following *C. elegans* strains were utilized for this work:

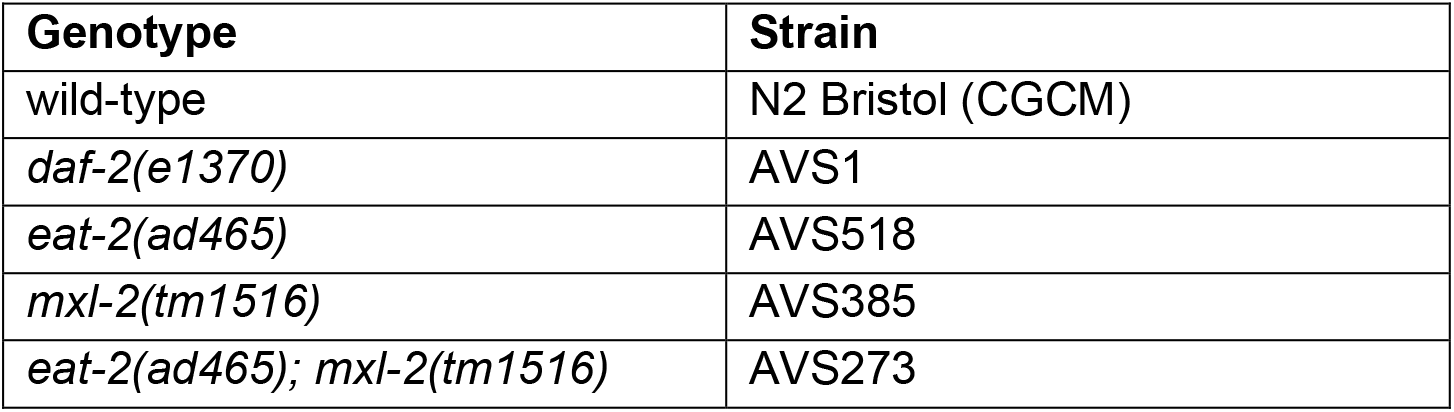

### C. elegans culture

Nematodes were maintained using standard laboratory culture techniques (Brenner and Brenner 1974). Animals were normally maintained on plates with *E. coli* OP50-1 as a food source. Animals for assays were raised on the RNAi-compatible *E. coli* strain HT115(DE3), transformed with empty vector (EV) plasmid or plasmid with an insert sequence complementary to the given gene of interest for feeding-based RNAi (Timmons and Fire 1998; Kamath et al. 2000). For assay plates, RNAi clones were grown overnight with shaking in LB and seeded onto agar plates containing 5 mM isopropylthiogalatoside (IPTG) and allowed to induce dsRNA expression overnight at room temperature; additional details can be found in (Cornwell and Samuelson 2020).

### Animal preparation and RNA isolation for RNA-Sequencing

Plates prepared for the RNA-Seq experiment were 10 cm RNAi plates seeded with 1 mL of 20x concentrated bacteria from an overnight culture. Plates were allowed to dry for 1-2 days at room temperature, also providing time for the induction of dsRNA production before adding animals. Approximately 3,000 L1 animals were then added, split across multiple plates to prevent starvation and overcrowding, and kept at 16 °C until the L4 stage. When animals reached L4, 600 µL of 4 mg/mL 5-fluorodeoxyuridine (FUdR) was added to each plate to prevent production of progeny, and the plates were transferred to 20 °C (“day 0”). Developmental stage was assessed separately for each strain to account for shifts in developmental timing.

At adult day 2, animals were collected in M9, washed twice in M9, and a final wash with DEPC-treated RNase-free water. Approximately two times the volume of the animal pellet of TRIzol® reagent was added to each animal preparation, followed by brief mixing and freezing overnight at -80 °C. Tubes were then allowed to partially thaw, and vortexed five minutes to assist with disrupting the cuticle. Each prep was transferred to a new tube, and 200 µL of chloroform per 1 mL of TRIzol was added, followed by vortexing for 20 seconds, and allowed to settle at room temperature for 10 minutes. Supernatants were transferred to new tubes, and an equal amount of 100% ethanol was added and mixed before proceeding to column purification with the Qiagen RNAeasy Mini kit (catalog 74104) according to the manufacturer’s instructions. Samples were eluted with 30 – 50 µL of DEPC-treated water and checked for initial concentration and quality with a Nanodrop ND-1000.

### RNA-Sequencing

Isolated RNA was provided to the University of Rochester Genomics Research Center for library preparation and sequencing. Prior to library preparation, an Agilent 2100 Bioanalyzer was used to confirm the RNA integrity of all samples. Libraries for sequencing were prepared with the Illumina TruSeq Stranded mRNA kit, according to the manufacturer’s instructions. Quality of the resulting libraries was checked again with a Bioanalyzer prior to sequencing to ensure integrity of the input material. Sequencing was performed on an Illumina HiSeq2500 v4, yielding an average of approximately 31 million single-ended 100 bp reads per sample. Quality of the output was summarized with FastQC (Andrews 2010) and reads were trimmed and filtered with Trimmomatic (Bolger et al. 2014). After filtering out low-quality reads, an average of 30.2 million reads per sample remained, and were used for the rest of the analysis.

### RNA-Sequencing data pre-processing

Samples were aligned to the *C. elegans* genome assembly WBcel235 with STAR 2.4.2a (Dobin et al. 2013) using gene annotation from Ensembl (version 82) (Cunningham et al. 2022). An average of 94.7% of reads per sample were uniquely mapped to the genome. These alignments were used as input to RSEM (version 1.2.23) for transcript-level quantification (Li and Dewey 2011) and featureCounts (version 1.4.6-p5) for gene-level counts (Liao et al. 2014). Ambiguous or multi-mapping reads, comprising an average of approximately 5% of the reads per sample, were not included in the gene-level count results.

Downstream analyses were performed using custom scripts run in R 4.0.2 (R Core Team). Gene identifiers were updated from the Ensembl annotation used for mapping (version 82) to version 106 for further analysis. Genes with little evidence of expression were removed prior to analysis of the count-based data by filtering out genes with fewer than 10 reads in every sample. Removing the unexpressed genes left 17,907 genes across five gene types considered (16,883 protein coding, 794 pseudogenes, 136 ncRNA, 87 lincRNA, and 7 snRNA). For exploratory analysis of expression trends across samples or genes, expression values normalized for library size were utilized, either TPM from RSEM or VST-normalized counts from DESeq2, as indicated.

### Differential expression analysis

Differential expression analysis was performed with DESeq2 1.30.1 (Love et al. 2014) in R, for all combination of strain and treatment against WT animals with empty-vector (EV) RNAi. Foldchange shrinkage procedures were applied to moderate fold-changes of low-expression genes. We considered significantly differentially expressed genes to be those with an FDR-corrected p-value < 0.05 and a foldchange magnitude of at least 2-fold (|log_2_ fold-change| ≥ 1). Fold-changes are for comparisons against an N2 reference population, unless otherwise specified. In a few instances, p-value results were smaller than what R can output in the default configuration, these are reported here in supplemental tables as 2.225074e-308, the smallest non-zero floating point number supported.

### Transcription factor target prediction

A workflow for prediction of transcription factor binding sites from positional weight matrix (PWM) motifs for *C. elegans* transcription factors was adapted from TargetOrtho 2 (Glenwinkel et al. 2021). Motifs were curated from CIS-BP (313), JASPAR (10), UniProbe (12), and manually from publications (7) for coverage of over one-third of known *C. elegans* TFs (Shih et al. 1995; Hume et al. 2015; Fuxman Bass et al. 2016; Allevato et al. 2017; Castro-Mondragon et al. 2022). Of motifs from CIS-BP, 202 are based on direct evidence from binding experiments (ChIP-Seq, PBM, etc) and the remainder are inferred by close homology to a TF in another species with direct experimental evidence. Motifs from JASPAR and UniProbe are based on experimental binding profiles. FIMO (Grant et al. 2011) was used to scan for motif matches in the *C. elegans* genome and seven other nematodes with available genomes: *C. brenneri*, *C. briggsae*, *C. remanei*, *C. japonica*, *P. pacificus*, *P. exspectatus*, and *A. lumbricoides*. TargetOrtho scripts were used to execute FIMO and pre-process the output. Initial identification of predicted binding within promoter regions was based on a symmetrical threshold distance from transcriptional start sites (TSS) of - 700/+700, and later filtered to -700/+100, within the range used for binding prediction to promoter regions in previous studies (Saito et al. 2013; Tepper et al. 2013). For FIMO match specificity, predicted associations with p < 0.0001 were considered, except for two of the manually curated E-box motifs, which had minimum p-values of 0.000195 and 0.00016; p < 0.0002 was used for these cases, as FIMO p-values are biased against short motifs. Results were read into R and, after filtering for the aforementioned -700/+100 TSS distance, two sets of motif matches were produced: 1) all *C. elegans* matches without further filtering, and 2) matches found in *C. elegans* and in the same region (upstream or downstream of the TSS) of a homologous gene in two or more of the other nematode species genomes; these are referred to as “all *C. elegans* motif matches” and “species homology-filtered motif matches”, respectively.

### Pathway and gene set enrichment

Gene sets for functional enrichment analysis were curated from the Gene Ontology, KEGG, Reactome, WormBase, WormExp, DIANA microT-CDS (miRNA binding predictions), and our TF target prediction set (Harris et al. 2009; Kanehisa et al. 2010; Blake et al. 2013; Paraskevopoulou et al. 2013; Yang et al. 2016; Gillespie et al. 2022). The R package GOSeq (1.42.0) was utilized for batch enrichment across many genes (Young et al. 2010). In some cases, over-representation analysis was performed for a smaller subset of gene sets of interest using Fisher’s exact test (hypergeometric test) either through the R package GeneOverlap (1.34.0) or the R function *phyper*.

### Analysis of publicly available datasets

Gene count results from multiple public datasets were re-analyzed in the same manner as our own dataset for differential expression. The public datasets and samples considered include *eat-2* and N2 animals on OP50 or HT115 bacteria at multiple timepoints in SRA SRP089617 (Tabrez et al. 2017); and *eat-2* and N2 animals at the L4 stage in GEO GSE125718 (Chen and Zou 2019). All differential expression analyses were only performed for samples within each dataset.

### Oxygen consumption

Animals were synchronized by bleach prep and added to RNAi plates with HT115 EV bacteria as a food source. FUdR was added at the L4 stage for each strain to prevent production of embryos, and animals were kept at 20°C. O_2_ consumption was assayed for whole animals at day 2 of adulthood with a Clark-type O_2_ electrode (S1 electrode disc, DW2/2 electrode chamber, and Oxy-Lab control unit, Hansatech Instruments, Norfolk UK). Animals were collected from plates in M9 buffer, pelleted briefly by centrifugation, and added to the electrode chamber in 0.5 mL of M9. After measurement of baseline respiration rate, FCCP (160 μM final concentration) was added to obtain maximal respiration measurements, followed by sodium azide (40 mM final concentration) for consumption rate with inhibited respiration. Each measurement continued until readings were stable, or up to 10 minutes. Post-measurement, animals were collected and frozen for protein quantification by Bradford assay to normalize oxygen consumption rates by total protein.

### Lifespan

For lifespan experiments, animals were kept from the L1 to L4 stage at 16°C, at which point FUdR was added to a final concentration of 400 μM; animals were then transferred to 20°C. Replica Set lifespan experiments were performed as detailed in (Samuelson et al. 2007; Cornwell et al. 2018, 2022). Briefly, in Replica Set experiments, individual animals are not followed over time, but rather a large population is split across 24-well “replica” plates, and a new sub-population of animals is scored at each time point and then discarded. Enough “replica” plates are set up at the outset of the experiment for each planned observation based on typical lifespan characteristics of the strains already established. Resulting data was fit to logistic function curve for determination of median lifespan, as previously described (Cornwell et al. 2018, 2022).

### Oxidative stress survival

Oxidative stress survival was performed similarly to (Johnson et al. 2014) but with observation and scoring facilitated by our version of the *C. elegans* Lifespan Machine (Stroustrup et al. 2014). Briefly, synchronized L1 animals were raised on HT115 bacteria with empty vector (EV) RNAi at 15 °C from L1 to L4 on 50 mm plates (Corning Falcon 351006). When animals reached the L4 stage, FUdR was added (400 μM final concentration) and then kept at 20 °C for the remainder of the experiment. When animals reached day 2 of adulthood, 7.7 mM *tert-*butylhydroperoxide (tBOOH) was added to the plates, and they were transferred to the lifespan machine scanners, with the plate lids in place, for imaging and analysis. The scan schedule was set such that every animal was analyzed approximately once per hour.

### Brood size

For each strain, animals were synchronized, and parent hermaphrodites were picked to their own plate at L4. Starting from 24 hours after L4, parents were picked to a new plate every 24 hours until no further eggs were observed. Larval animals were counted after 48 hours, allowing enough time for all viable eggs to hatch. Eggs remaining unhatched after this time were considered dead and recorded. Animals were kept at 20 °C on HT115 EV *E. coli* as a food source. Two independent trials were conducted. Pairwise t-tests were performed to assess the statistical significance of differences between conditions.

## Results

### DR induces an adaptive transcriptional response to a chronic reduction in food availability

We sought to identify changes in gene expression associated with dietary restriction, with and without perturbation of the MML-1::MXL-2 complex via the *mxl-2(tm1516)* null mutation. We conducted transcriptome-wide RNA-sequencing of chronologically age-matched day 2 adult animals, including: wild-type (N2), *eat-2(ad465),* and *mxl-2(tm1516)* single mutant animals, as well as *eat-2;mxl-2* double mutant animals. Additionally, we included loss of *pha-4* (ortholog of FoxA, a forkhead transcription factor) a modulator of gene expression in response to nutrient stress (Zhong et al. 2010; Pandit et al. 2014), required for the increased lifespan of *eat-2* mutant animals (Panowski et al. 2007). Furthermore, we included loss of *daf-16* (FoxO) as a putative negative control, as *daf-16* has been shown to be dispensable for the increased lifespan of *eat-2* mutant animals, but is a central regulator of longevity in other contexts (Lakowski and Hekimi 1998; Tepper et al. 2013).

We first assessed the overall quality and fidelity of the gene expression for each condition. We were able to reliably assess gene expression for 17,907 genes, after removing consistently unexpressed genes and gene types without evidence of polyadenylation. We conducted a principal component analysis (PCA) across all 17,907 genes and found at least one principal component could account for substantial variability in the overall dataset in which biological replicate samples cluster together (Figure 1A). The Pearson *r* for pairwise correlations among intra-group biological replicates is higher than 0.9 for all but two groups, and not lower than 0.85 in any case, implying reliably consistent patterns of gene expression between biological trials (Figure S1A). We find that the variability between biological replicates is associated with PC2 for *eat-2(ad465)* controls and *eat-2(ad465);daf-16(RNAi)*; PC1 is associated with *mxl-2(tm1516)* control RNAi, *daf-16(RNAi)*, and *pha-4(RNAi)*. We also observed that this variability is not associated with known batch variables (Figure S1B). As expected, we found inactivation of *daf-16* in *eat-2* animals had very little effect on gene expression, as fold-changes between *eat-2* controls and *eat-2 daf-16(RNAi)* are highly correlated (Pearson r = 0.92, Figure 1B (i)), consistent with conclusions based on genetic interactions. In contrast, loss of either *mxl-2* or *pha-4* dramatically disrupted the gene expression profile of *eat-*2 mutant animals (Figure 1B (ii) and (iii) respectively). The impact of *mxl-2* and *pha-4* loss on *eat-2* gene expression aligns with previous genetic analysis: *pha-4* and *mxl-2* are required for *eat-*2-mediated increases in longevity (Panowski et al. 2007; Johnson et al. 2014). Collectively, we conclude our gene expression profiles have high fidelity and consistency, and genetic perturbation of *mxl-2*, *pha-4*, and *daf-16* alters gene expression of *eat-2* mutant animals as predicted based on previously characterized genetic interactions.

**Figure 1.**
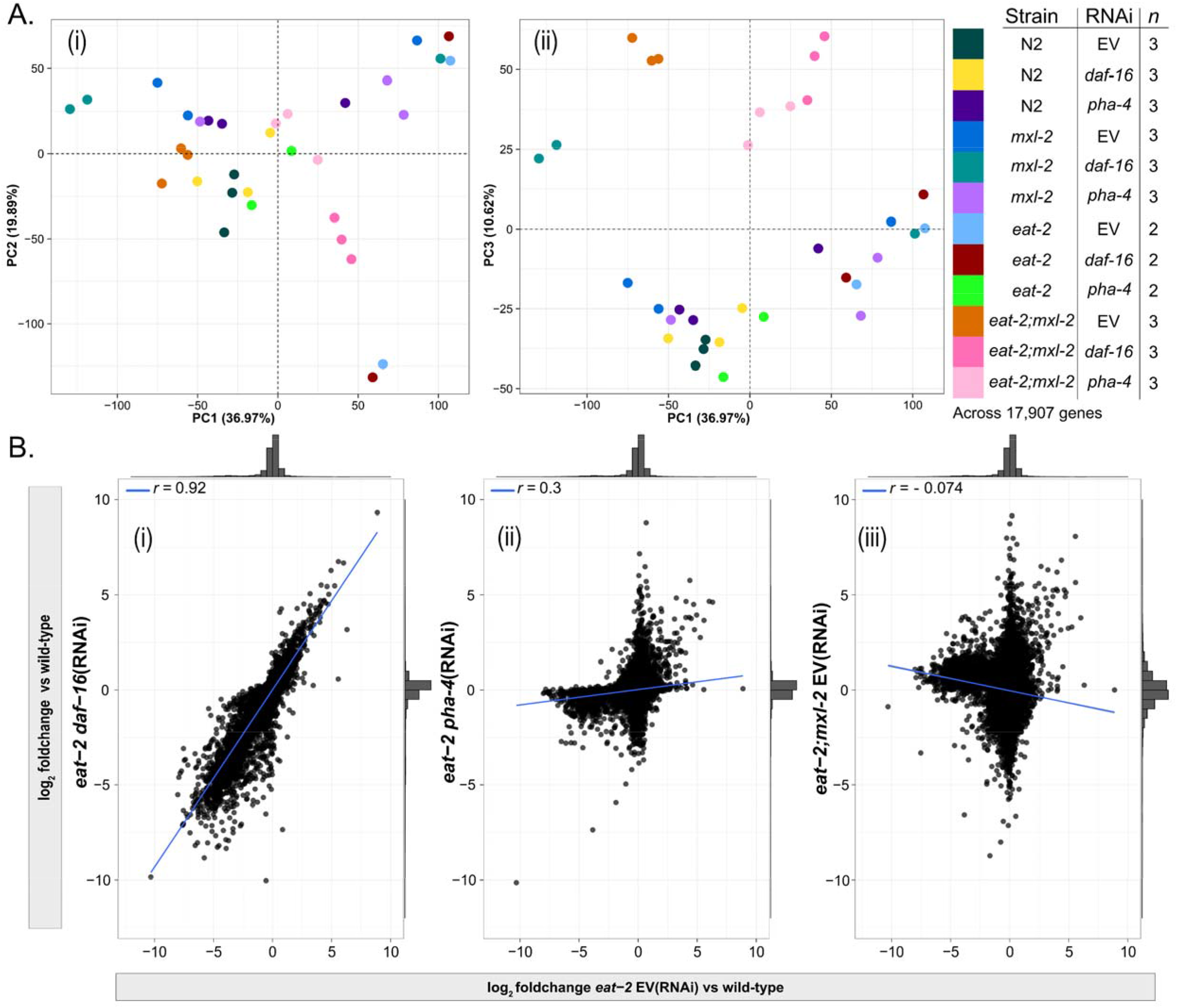
Exploratory analysis across all expressed genes shows relative gene expression changes in *eat-2* are broadly disrupted by loss of *pha-4* and *mxl-2* but not *daf-16*, in accordance with results of genetic experiments. **(A)**. At least two biological replicates were analyzed across 17,907 genes expressed in any sample. Principal component analysis on variance-stabilized (VST) normalized count data shows that there are both distinct clusters between some sample groups, and that biological replicate samples cluster together in at least one principal component that explains a substantial proportion of the variance of the overall dataset. The Pearson *r* for pairwise correlations of VST gene expression between replicate samples across all expressed genes was not lower than 0.85 in any case (Figure S1A). We observed that variability between biological replicates was not associated with the batch in which the samples were prepared (Figure S1B). The key on the right shows the correspondence between color and sample group (strain, RNAi treatment) and the number of biological replicate samples for that group. **(B)**. Comparison of relative expression changes (log_2_ fold-change) across all 17,907 genes expressed in any sample in *eat-2* DR animals relative to WT (x-axis) and *eat-2* with *daf-16* RNAi (i), *pha-4* RNAi (ii), or *eat-2;mxl-2* double mutant animals (iii), each also relative to WT (y-axis). Wild-type is N2 with empty vector RNAi. Similar to results of genetic analysis with lifespan phenotypes, *daf-16* is not required for maintenance of the *eat-2* DR gene signature (i), but loss of either *pha-4 (*ii) or *mxl-2* (iii) result in dramatic shifts in gene expression. Pearson correlation indicates a strong and significant positive linear association between *eat-2* vs WT and *eat-2;daf-16* (RNAi) vs WT (r = 0.92, p < 2.2×10^-16^), but a much weaker positive trend after loss of *pha-4* (r = 0.3, p < 2.2×10^-16^), and a weak negative association after *mxl-2* loss (r = -0.074, p < 2.2×10^-16^). The histograms on the side of each panel indicate the density of points.

We sought to identify the transcriptional adaptive response to DR and compared the gene expression signature of *eat-2* mutant animals to wild-type (N2) animals through differential expression (DE) analysis. We found 1704 genes with significantly altered expression and at least a moderate fold-change in *eat-2* animals, of which 249 genes (14.6%) were up-regulated and the majority-1455 (85.4%)-were down-regulated (Figure 2A) (DESeq2 FDR-corrected p-value < 0.05, absolute log_2_ fold-change ≥ 1). Among DE genes in *eat-2* mutant animals, 75 had prior associations to lifespan phenotypes (based on Wormbase WS282 (Davis et al. 2022)): most in down-regulated genes, with more than half associated with extended longevity with loss of function, correlating down-regulation in long-lived DR animals. As about 15% of *C. elegans* genes are co-transcribed in operons (Blumenthal et al. 2002), we also looked for enrichment of annotated operons, and found enrichment of five operons corresponding to 11 genes among all differentially expressed genes in *eat-2*, indicating that regulation of transcription at the level of operons has little influence on the DR transcriptional profile. The complete set of differential expression results can be found in Table S1.

**Figure 2.**
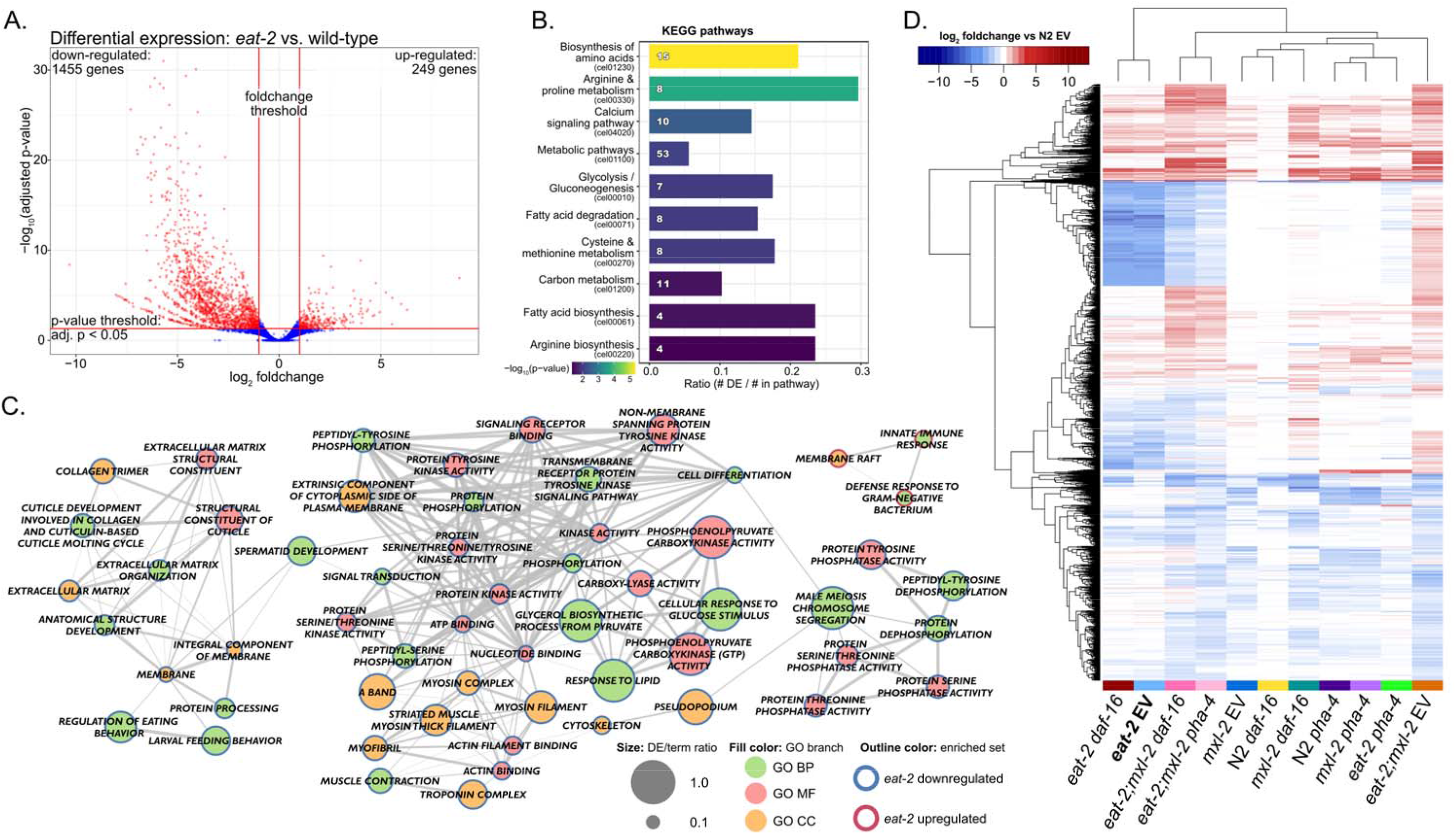
DR in *eat-2* animals significantly changes expression of more than 1000 genes, providing evidence of broad metabolic reprogramming. **(A.)** Volcano plot demonstrating the strategy for differential gene expression analysis. The x-axis shows log_2_ fold-changes for *eat-2* EV relative to wild-type (N2 EV), and the y-axis shows -log_10_ transformed FDR-adjusted p-values from DESeq2 for the same comparison. Vertical red lines indicate the threshold used for fold-change magnitude (|log_2_ FC| ≤ 1) and horizontal red line indicates the p-value threshold (adjusted p-value < 0.05). With these criteria, 1,455 genes were significantly down-regulated and 249 were upregulated in *eat-2* (red points). The same criteria were also applied to all other comparisons. **(B.)** KEGG pathways significantly enriched in *eat-2* down-regulated genes indicate changes in multiple aspects of metabolism, including amino acid biosynthesis, fatty acid metabolism, and energy-associated pathways. The bar length is the fraction of the pathway genes that overlap with the *eat-2* gene set, the number in the bar is the number of overlapping genes, and bar color shows the −log_10_ adjusted p-values (GOSeq). All pathways shown had significant enrichment (adjusted p-value < 0.05). **(C.)** Significantly enriched Gene Ontology (GO) terms for genes differentially expressed with DR in *eat-2*. Network representation allows visualization of related terms which share member genes. Up- and down-regulated genes were treated as separate gene sets (red and blue node outline, respectively). Node fill color indicates the GO sub-ontology a term belongs to: biological process (BP), molecular function (MF), or cellular component (CC). Node size indicates the proportion of genes associated with the term that were present in the gene set. Connections between nodes (edges) show terms with shared member genes, with thicker edges representing a greater degree of overlap. **(D.)** Heatmap of log_2_ fold-changes across the 4,762 genes differentially expressed (DE) in at least two of any of the indicated comparisons relative to wild-type. Non-significant fold-changes may also be indicated for a given comparison if the gene was significant in other comparisons.

In order to identify functionally how expression changes may contribute toward the aging benefits of DR, we performed over-representation analysis to identify significant associations with pathways and gene sets. Among the genes downregulated in *eat-2* mutant animals, metabolic reprogramming, including reduced expression of genes involved with amino acid and fatty acid biosynthesis were significantly enriched. In contrast, analysis of *eat-2* upregulated genes produced no significantly enriched KEGG or Reactome pathways, indicating a lack of broadly coordinated activation of gene expression across the organism in response to DR (Figure 2B). Macromolecule metabolites have been previously observed to be less abundant in *eat-2* through metabolomics experiments (Mouchiroud et al. 2015; Gao et al. 2018) and overall protein content is lower in *eat-2* animals (Yuan et al. 2012; Gao et al. 2018), which is consistent with our findings. We also found downregulated expression of genes regulating fatty acid metabolism; *eat-2* mutant animals are thin with reduced fat stores (Avery 1993; Heestand et al. 2013; Bar et al. 2016). A switch to fatty acids as an energy source is important in DR (Heestand et al. 2013; Weir et al. 2017), and *eat-2* mutant animals have largely depleted reserves of lipids. Interestingly, *elo-5*, a fatty acid elongase involved in monomethyl branched-chain fatty acid synthesis, which is important for lifespan of wild-type and *daf-2* animals, as well as glucose-induced stress resistance, is significantly upregulated in our *eat-2* samples, indicating possible adaptive upregulation of specific fatty acids to maintain membrane integrity in the context of low overall lipid availability (Kniazeva et al. 2004; Samuelson et al. 2007; Vieira et al. 2022)(Table S1). We conclude *eat-2* mutant animals globally downregulate significant components of metabolic gene expression to match the diminished availability of nutrients and optimize energy utilization, which suggests that expression levels of metabolic genes are causally linked with nutrient availability.

### Metabolic remodeling during DR re-allocates energy away from costly processes toward somatic health via changes in gene expression

We next assessed enrichment of broader functional terms from the Gene Ontology in *eat-2* animals. We found the only terms enriched for *eat-2* upregulated genes are associated with innate immune and bacterial defense responses (Figure 2C), however we also noted that the upregulation of a small set of innate immune response genes was not specific to *eat-2* and also occurred with *mxl-2* mutation or *pha-4* RNAi under both basal conditions and in *eat-2* (Figure S2). The lack of concerted regulation of functions associated with *eat-2* upregulated genes suggests that chronic DR is not activating transcription as part of a specific adaptive response to nutrient stress by day two of adulthood.

Rather, we observed reduced gene expression associated with several biological processes consistent with a shift from reproduction to survival. For example, there is significant enrichment for downregulation of sperm-associated gene sets, including: spermatid development, sperm meiosis chromosome segregation, sperm motility via pseudopodia, and Major Sperm Proteins (MSPs, 31 of 47). MSPs are the most abundant proteins in *C. elegans* sperm and necessary for reproduction, both as structural components and signals for oocyte maturation (Harris et al. 2006; Shimabukuro and Roberts 2013), (Figure 2C). We posit this represents an adaptive shift away from reproductive fecundity in favor of survival. The total number of sperm within the hermaphrodite spermatheca limits the progeny number in unmated hermaphrodites, which typically form prior to the onset of adulthood. *eat-2* animals have smaller total brood sizes and extended reproductive-span (Crawford et al. 2007; Albert Hubbard 2007; Luo et al. 2009; Korta et al. 2012). Our results suggest down regulation of gene expression associated with sperm production provides a simple but effective mechanism for animals undergoing chronic DR to prioritize survival over investing in the energetically costly process of producing progeny; this is in contrast to the response to acute starvation in reproductive adult wild-type animals in which animals retain eggs, leading to internal hatching and matricide to promote survival of progeny (Chen and Caswell-Chen 2004).

We found a significant number of genes related to muscular function down-regulated. For example, decreased expression of genes associated with the A band of the sarcomere and the myosin complex (Figure 2C), which is consistent with thinning of body wall muscles observed in *eat-2* mutant animals (Mckay et al. 2004). Additional genes include those broadly-expressed in the body wall-muscle, including: *unc-54, myo-3* as well as *mlc-3* (log_2_ foldchange -1.32 to -2.3), which encode myosin heavy chain and light chain proteins respectively (Benian and Epstein 2011). We posit downregulation of muscle gene expression results in the maintenance of a smaller volume of muscle, thereby limiting the drain of energy reserves from functions vital to maintaining viability. Concurrently, protein from skeletal muscle provides energy during severe nutrient restriction (Sharples et al. 2015).

Increased expression collagens has been associated with longevity, particularly under reduced insulin-like signaling in *daf-2* animals (Ewald et al. 2015; Sandhu et al. 2021; Rahimi et al. 2022). However, we found collagen-associated genes significantly down-regulated in *eat-2* mutant animals (e.g., collagen trimer, constituent of cuticle, ECM constituent). Despite this, *eat-2* animals do not exhibit compromised cuticle integrity, and retain properties of “young” cuticle into old age (Essmann et al. 2020). Decreased expression of collagen genes is also consistent across analysis of public *eat-2* gene expression datasets in late-larval animals at L4 (Chen and Zou 2019), and young adults at day 1 and 3 (Tabrez et al. 2017). Downregulation of collagens has been observed in mice under DR, suggesting an evolutionarily conserved mechanism (Choi 2020). As collagen is the most abundant protein in the *C. elegans* cuticle and covers the entire outside of the animal (Sandhu et al. 2021), we posit synthesis of new collagen to be highly regulated in *eat-2* due to restricted energy availability. Collagen usually consists of trimeric repeats of glycine and two variable amino acids-often proline and hydroxyproline (Cox et al. 1981; Johnstone 2000). While these are not among the most bioenergetically costly to synthesize, and are also not among the diet-derived essential amino acids in *C. elegans,* the total amount of collagen in the cuticle and ECM still makes collagen maintenance an energetic burden (Akashi and Gojobori 2002; Zečić et al. 2019; Sandhu et al. 2021). Interestingly, supplementing WT animals with low concentrations of proline or glycine is sufficient to extend lifespan but does not further increase lifespan of *eat-2(ad1116)* animals (Edwards et al. 2015). Overall, the consistent downregulation of collagens across multiple *eat-2* datasets again suggests an energetic “cost-cutting” approach in DR, to direct limited resources to the most acute needs and provides an additional energy resource, though it remains unclear if DR activates other strategies to reduce unnecessary collagen turnover, or if collagen synthesis would be upregulated to heal wounds in DR animals subject to cuticle damage (Mesbahi et al. 2020).

A large number of genes with kinase or phosphatase activity are down-regulated in *eat-2* mutant animals-111 and 76, respectively (Figure S3A). To determine whether gene products were associated to specific signaling cascades, or enriched in specific tissues, we looked for pathways and functions associated with this gene subset. Looking at all associations, we found kinases associated with arginine metabolism (*F46H5.3, F32B5.1, W10C8.5*), glycolysis and gluconeogenesis (*pfk-1.2, pck-2, pck-1*) and phosphatases linked to mRNA surveillance (*C09H5.7, C34D4.2, F52H3.6, ZK938.1, gsp-4, gsp-3, T16G12.7*) (Figure S3B-C). Arginine kinases play a role in energy homeostasis by helping to meet acute increases in energy demands (Fraga et al. 2015); for example, overexpression *argk-1,* extends *C. elegans* lifespan through activation of AAK-2 (AMPK) (Curtis et al. 2006; McQuary et al. 2016). *C. elegans* possesses more than 25 phosphatases with substantial homology to yeast GLC7 (WormBase WS282), which is important for mRNA export from the nucleus-an aspect of mRNA surveillance (Gilbert and Guthrie 2004; Izaurralde 2004; Davis et al. 2022). Of the seven GLC7 homologs with reduced expression in *eat-2*, GSP-3/4 are the most well-studied, with roles in spermatogenesis that mirror the function of homologs expressed in male mice (Chu et al. 2006). More broadly, both the kinases and phosphatases down-regulated in *eat-2* mutant animals are enriched in functions related to the development of sperm (Figure S3D); kinase and phosphatase expression is a putative mechanism to regulate spermatid function without translation of new proteins through post-translational modulation of protein activity, as spermatids lack ribosomes (Reinke et al. 2000; Ellis and Stanfield 2014).

### MXL-2 regulates expression of a distinct set of genes during dietary restriction

We next sought to identify genes with altered levels of expression in our other sample groups. We proceeded to run differential expression analysis for single mutant genetic backgrounds: *eat-2(ad465), mxl-2(tm1516),* along with *eat-2(ad465);mxl-2(tm1516)* double mutants after treatment with either control RNAi (empty vector), *pha-*4*(RNAi)*, or *daf-16(RNAi)*. These genetic perturbations were compared to wild-type animals treated with empty vector RNAi in a pairwise manner, using the same criteria for significant expression changes (Table 1). Across these 11 contrasts, a total of 4762 genes were differentially expressed relative to N2 EV in at least two comparisons (Figure 2D). Loss of *daf-16* has little effect on the transcriptional response to chronic DR, as expected. Also, in accordance with genetic analysis of requirements for *eat-*2 lifespan, loss of *pha-4* effectively negated the gene expression signature of *eat-2* animals: the *eat-2;pha-4* profile more closely resembles *pha-4* inactivation in wild-type and *mxl-2* mutant animals (Figure 2D).

**Table 1.**
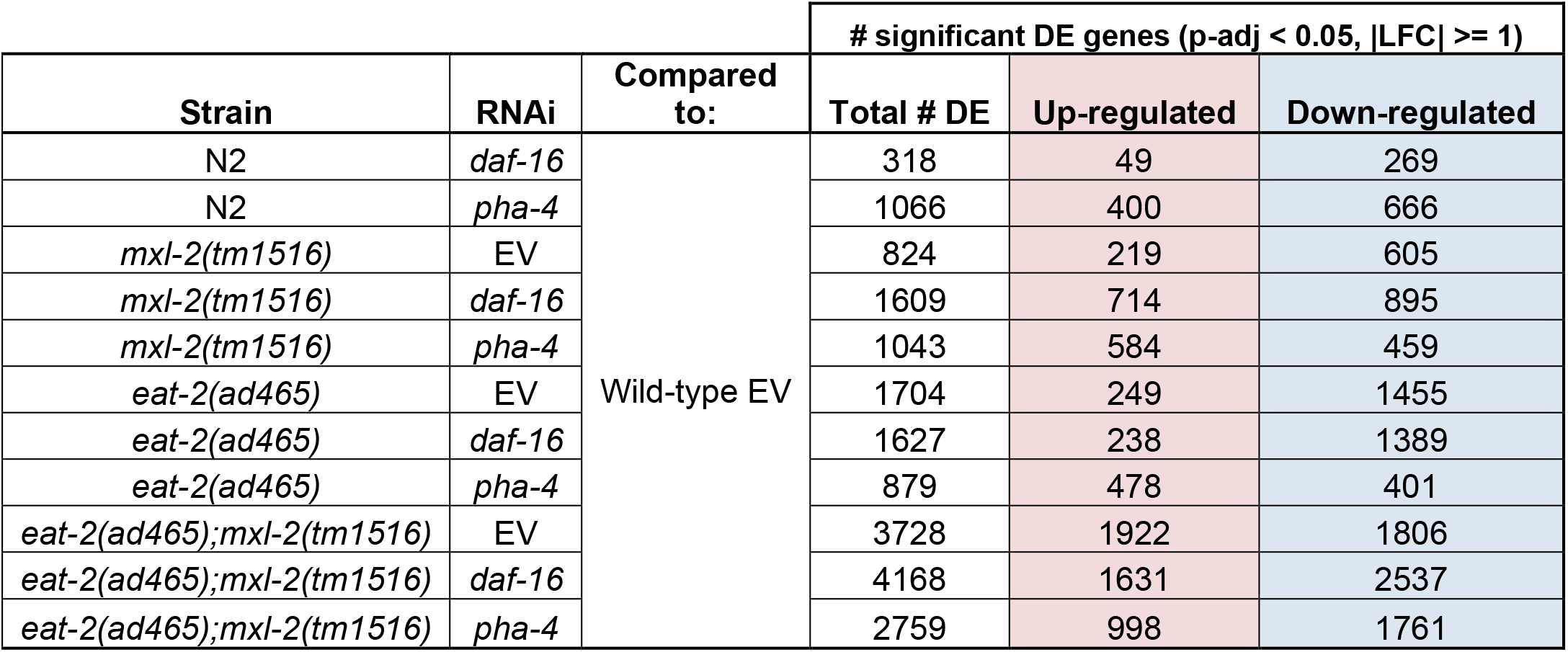
Summary of significantly differentially expressed (DE) genes for all considered samples compared to wild-type EV. Summary of differential expression analysis for genes with significant changes in expression for all considered samples compared to wild-type (N2 EV). Significant DE genes were based on the following criteria after analysis with DESeq2: adjusted p-value < 0.05 and log_2_ fold-change magnitude ≥ 1. Up-regulated genes exhibited higher expression (positive fold-change) in the subject condition compared to N2 EV, and vice-versa for down-regulated genes (negative fold change).

Comparison of the *mxl-2(tm1516)* null mutation alone to wild-type animals results in significant expression changes to more than 800 genes, with 75% of those down-regulated; *mxl-2* loss was distinct from that of either *pha-4* or *daf-16*, the latter of which yielded the fewest total DE genes of any of these comparisons (318 total, 49 up and 269 down), which may reflect typically minimal DAF-16 activity under basal conditions (Shaw et al. 2007). In contrast, loss of *mxl-2* drastically changed the gene expression pattern of *eat-2* mutant animals. Surprisingly, inactivation of either *pha-4* or *daf-16* in *eat-2;mxl-2* mutant animals yields profiles distinct from both *eat-2*;*pha-4* and *eat-2;mxl-2*; see section on synthetic dysregulation further on for additional related results and discussion.

We looked for tissue-associated biases in differential gene expression, based on sets of genes enriched in muscle, neurons, hypodermis, intestine (Kaletsky et al. 2018), or in sperm (Reinke et al. 2000) (Figure S4A). In agreement with our Gene Ontology analysis of *eat-2* expression changes, we found a substantial and significant set of downregulated genes in *eat-2* mutant animals to be enriched in sperm, as well as significant over-representation of muscle-associated genes. The intestine is commonly referred to as the most metabolically active tissue in *C. elegans*, and both the MML-1::MXL-2 complex and PHA-4 act in intestinal cells in adult animals (Panowski et al. 2007; Johnson et al. 2014; Nakamura et al. 2016; Horowitz et al. 2023); we found enrichment for intestinally expressed genes in both the up- and down-regulated transcriptional response to *eat-2*, *mxl-2*, or *pha-4* single-perturbation conditions. Intriguingly, we found enrichment of neuron-associated genes among those down-regulated in *mxl-2* null mutant animals, which suggests Myc-family TFs may coordinate organismal adaptive responses from the nervous system.

### MXL-2 and PHA-4 are required for the majority of gene expression reprogramming associated with chronic dietary restriction

We next identified the portion of the *eat-2* expression signature that required *mxl-2* and/or *pha-4*. We defined *mxl-2-*dependent genes as: 1) differentially expressed in *eat-2*, 2) not *similarly* differentially expressed in *eat-2;mxl-2* (i.e., not changing in the same direction), and 3) specific to the DR background (i.e., unchanged in *mxl-2*) -all compared to wild-type. This strategy was applied to both *eat-2* up-regulated and down-regulated genes (Figure 3A i,ii), for dependence on *mxl-2, pha-4,* and *daf-16* (Figure 3B, Supplementary Table 2). 89% of genes down-regulated in *eat-2* mutant animals required *mxl-2* and 92% required *pha-4*, while only 19% required *daf-16*. The pattern is similar for genes up-regulated in *eat-2*, but less pronounced, with 54% of *eat-2* upregulated genes requiring *mxl-2*, 65% requiring *pha-4*, and only 31% requiring *daf-16*. Overall, the vast majority of genes down-regulated genes in *eat-2* require both *pha-4* and *mxl-2*, but not *daf-16*, to maintain transcriptional suppression (Figure 3C).

**Figure 3.**
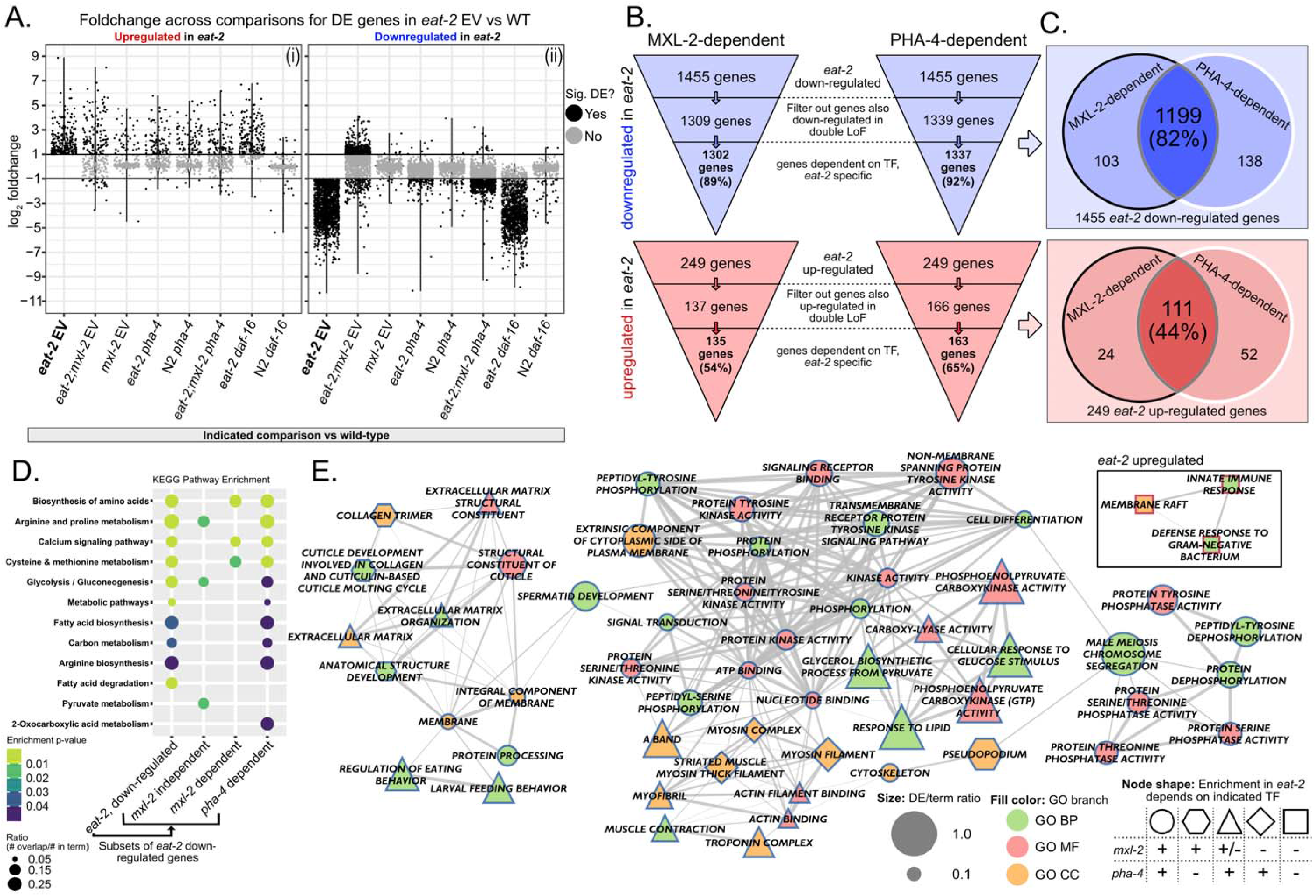
Both *mxl-2* and *pha-4* are required for the majority of the DR gene signature in *eat-2* animals. **(A.)** DR genes dependent on *mxl-2* were those up (i) or down (ii) regulated in *eat-2* that were not significantly regulated in a similar manner in *eat-2;mxl-2* animals. Genes with significant and opposite fold-change in *mxl-2* single mutant animals were filtered out for specificity to the DR (*eat-2*) context. The same strategy was also applied for investigating *pha-4*-dependent genes. The results are illustrated in (B) and Table S2. For each sub-panel A (i) and (ii), the genes represented in each column are the same, but the fold-changes and differential expression calls are for the indicated comparison. Horizontal lines at 1 and -1 indicate the foldchange threshold. The horizontal spread of points is random jitter added to reduce overplotting. **(B.)** Schematic representation of the filter steps to determine the *eat-2* genes dependent on *mxl-2* or *pha-4*. We found that a slightly larger proportion of *the eat-2* signature depends on *pha-4*, but the vast majority of the downregulated portion of the signature require both **(C)**. **(D, E)** Pathway (D) and Gene Ontology (E) enrichment for TF-dependent *eat-2* genes. Over-representation analysis was run on each list independently. Genes from a term that was significantly enriched in *eat-2* may be split between “dependent” and “independent” which can result in the term not showing as significantly enriched in either category (e.g. “Fatty acid biosynthesis” in (D)). (E) shows the same network representation for *eat-2* GO term enrichment as Figure 2C, with node shapes indicating TF dependency (see legend on bottom-right). As expected based on (B) and (C), many pathways and functions are dependent on both factors, with an overall greater proportion dependent on *pha-4*. Notably, some muscle-associated terms (mysoin filament, myosin complex) are *mxl-2*-independent but *pha-4*-dependent, and vice-versa for some collagen and sperm-associated terms (pseudopodium and collagen trimer, respectively). Triangle-shaped nodes are dependent on *pha-4* but lacked significant enrichment in both the *mxl-2*-dependent and -independent sets. The only GO terms with significant enrichment in the DR upregulated genes are independent of both factors, and a subset of these genes are upregulated with both loss of *mxl-2* and *pha-4* outside of the context of DR, indicating they may be part of a non-specific low-level stress response (see Figure S2).

We conducted functional enrichment analysis with the *eat-2* gene sets that were either dependent or independent of *mxl-2* or *pha-4* (Figure 3D-E). No KEGG pathways were significantly enriched in *pha-4*-independent genes. In contrast, the glycolysis, gluconeogenesis, as well as arginine and proline metabolism pathways were *mxl-2* independent (Figure 3D). We noted that the four *mxl-*2-independent *eat-2* downregulated genes in arginine/proline metabolism function in catabolism of these amino acids; while proline catabolism has been linked to longevity in other contexts (Zarse et al. 2012; Pang et al. 2014), down-regulation in *eat-2* may be concomitant with downregulation of collagen synthesis genes, as collagen is rich in proline/hydroxyproline (Cox et al. 1981; Johnstone 2000). Among functional annotations from the Gene Ontology (GO), genes of some muscle-related terms-myosin complex and myosin filament-are *mxl-2-*independent but *pha-4*-dependent, with the remaining muscle functions and components being dependent on *mxl-2* (Figure 3E). A few GO terms were independent of *pha-4* but dependent on *mxl-2*, including pseudopodium (associated with sperm development and motility) and collagen trimers. Sperm-enriched genes down-regulated in *eat-2* mutant animals are broadly dependent on *mxl-2* (Figure S4B). Taken together, we posit the de-repression of some of these energy intensive processes with loss of the *mxl-2* or *pha-4* transcription factors prevents the adaptive allocation of resources toward somatic maintenance, resulting in compromised health and lifespan.

### *C. elegans* Myc-family members possess unexpected transactivation domains

In *C. elegans*, the consensus to date is that the MML-1::MXL-2 complex activates and the MDL-1::MXL-1 complex represses transcription (Yuan et al. 1998; Pickett et al. 2007). Consistent with their different activities, we previously found loss of *mdl-1* or *mxl-1* increased longevity, while loss of *mml-1* or *mxl-2* decreased longevity, and that these genetic interactions are epistatic (Johnson et al. 2014). These results suggested that under otherwise basal conditions, the two complexes converge on shared target sites and compete to regulate longevity-associated gene expression. In agreement with the Y2H studies of MML-1::MXL-2 and MDL-1::MXL-1 complex activity (Pickett et al. 2007), mammalian Mondo proteins contain a transactivation domain (TAD) (Mcferrin and Atchley 2012; Conacci-Sorrell et al. 2014; Ceballos et al. 2021) and mammalian Mad complexes oppose the transcriptional activity of Myc (Lahoz et al. 1994; Schreiber-Agus et al. 1994; McFerrin and Atchley 2011). Thus, we were surprised to find *mxl-2* dependence for the downregulation of gene expression in *eat-2* animals; the clear prediction based on molecular annotation, known function, and genetic analysis suggested that the MML-1::MXL-2 complex would be necessary for activating transcription under conditions of DR. While the *mxl-2* dependence for the vast downregulation of gene expression in *eat-*2 could be indirect, we considered the possibility that Myc-family members may not unidirectionally regulate transcription in every context. Using recently developed neural-network models for TAD prediction, we applied these models to the *C. elegans* Myc-family members and their human orthologs (Erijman et al. 2020; Sanborn et al. 2021). As expected, we found a high-confidence predicted TAD in MML-1 (Figure S5A), within amino acid residues 245-276, which parallels the presence of the TAD in Mondo-conserved regions IV and V in vertebrates, and is also predicted by ADPred (Figure S5A) (Mcferrin and Atchley 2012; Ceballos et al. 2021). To our surprise, a high-confidence TAD was also predicted on the C-terminus of MDL-1 (Figure S5C). In contrast, human MXD/MAD family proteins show much weaker confidence of a predicted TAD in MXD1 and MXD3, which suggests in some instances MDL-1 may activate transcription, in contrast to the mammalian homologs. We also found potential TAD regions of lower confidence on the C-terminal side of MXL-2 and MXL-3, but not MXL-1 (Figure S5B, D). Intriguingly, the wider presence of TADs within *C. elegans* Myc family members may explain why *C. elegans* lack a true Myc-ortholog and loss of any particular Myc-family member still results in viable animals; yet loss of Myc is lethal in higher metazoans.

### Dysregulated gene expression in *eat-2;mxl-2* compromises reproductive fitness

Given the alterations in gene expression linked to reproduction in *eat-2* animals, we assessed the impact of *mxl-2* loss on *eat-2* fecundity. We quantified progeny production through reproductive lifespan (i.e., the period of active egg laying in reproductive hermaphrodites), and whether animals produced unhatched eggs. *eat-2* animals are known to have smaller brood sizes, and exhibit extended reproductive lifespan (Crawford et al. 2007). In the absence of *mxl-2*, *eat-2* mutants produce a significantly reduced total brood size— almost half that of *eat-2* alone— without affecting the duration of the egg-laying period (Figure 4A,B, Table S3). Furthermore, dead eggs were only observed from *eat-2;mxl-2* hermaphrodites (Figure 4C, Table S3). Interestingly, *pha-4* inactivation is known to reduce *eat-2* reproductive lifespan (Luo et al. 2009); in contrast, in the absence of *mxl-2* we find the duration of the reproductive lifespan of *eat-2* mutant animals is intact, but negatively impacts total fecundity. Given the limited nutrient resources of animals experiencing chronic DR, and our observation of de-repression of many metabolic and reproduction-associated genes in *eat-2;mxl-2* mutant animals, we posit that aberrant reallocation of resources in *eat-2;mxl-2* yields uneven provisioning to progeny and thus critically compromises the fitness of a portion of embryos (see Figure S6 and Table S3 for additional trial data).

**Figure 4.**
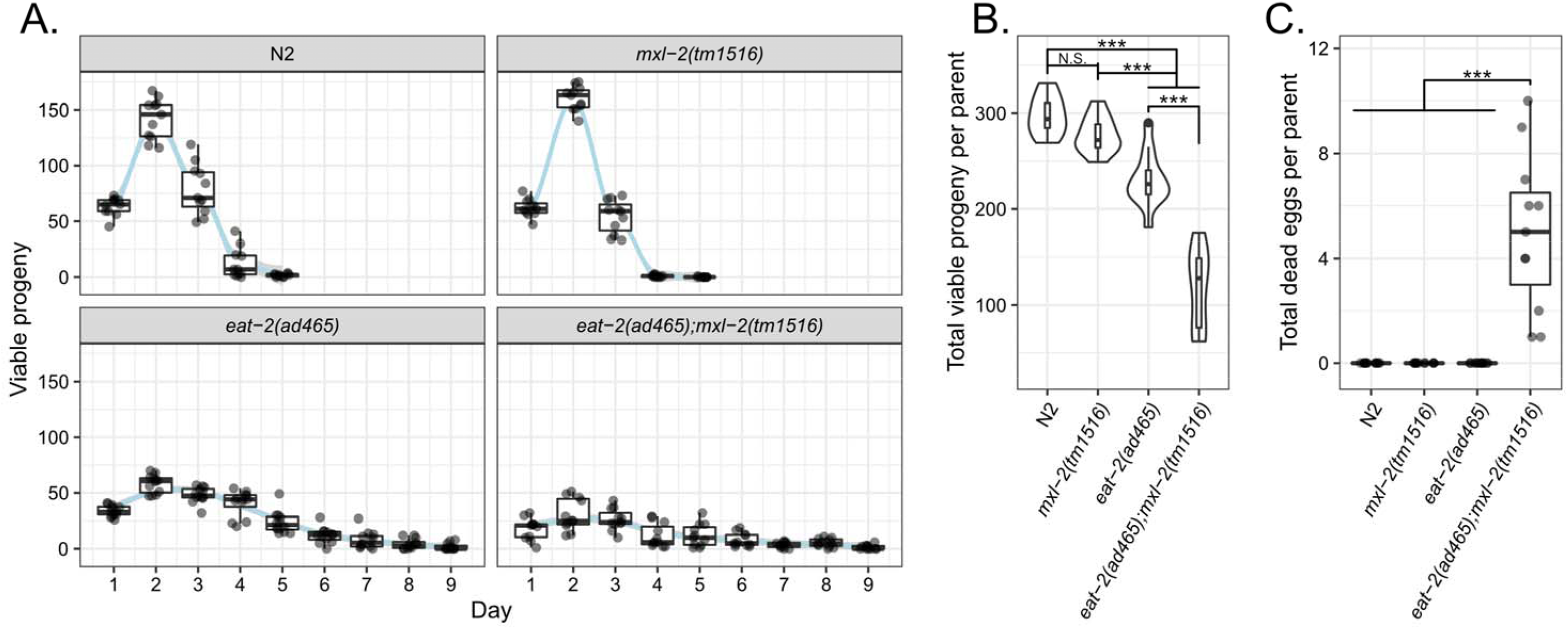
Loss of MXL-2 in DR but not ad libitum animals compromises fecundity and embryo viability. Brood size assays were performed for N2, mxl-2, eat-2, and eat-2;mxl-2 animals. For each strain, animals were synchronized and parent hermaphrodites were picked to their own plate at L4. Starting from 24 hours after L4, parents were picked to a new plate every 24 hours until no further eggs were observed. Larval animals were counted after 48 hours, allowing enough time for all viable eggs to hatch. Eggs remaining unhatched after this time were considered dead and recorded. Dietarily restricted animals are reproductive through a longer period of their life (”reproductive-span”), as previously reported (Crawford et al. 2007; Luo et al. 2009), which we find MXL-2 to not be required for (A). However, the smaller brood size typical of eat-2 animals was further reduced in eat-2;mxl-2 double mutants (B). Furthermore, dead eggs were observed for double mutants, but not the other strains (C). Data shown from a representative trial. See Table S3 for additional trial results and statistical analysis, and Figure S6 for the plotted additional trial results.

### Lower food intake in DR does not significantly alter respiratory rate, and is not affected by MXL-2 loss in young adult animals

We ascertained whether other physiological aspects of a DR-optimized metabolism were disrupted in the absence of *mxl-2*. Multiple groups have shown, perhaps counter-intuitively, that respiratory rates are as high or even higher in *eat-2* mutant animals than in wild-type (Houthoofd et al. 2002; Edwards et al. 2013; Lim et al. 2018). Surprisingly, we found no significant difference in baseline respiratory rate, maximal, or reserve respiratory capacity between N2, *mxl-2*, *eat-2*, or *eat-2;mxl-2* mutant animals (Figure 5A-C). Given that only a subset of the lipid metabolism genes that are part of our DR expression signature are dependent on *mxl-2*, a normal respiratory rate may be maintained by a switch to fatty acid oxidation in DR that does not require MXL-2. It remains to be determined if *pha-4* loss alters DR metabolic rate. Age may also play a role; recent work reported that *eat-2* animals maintain “youth-like” mitochondrial membrane polarity at more advanced age than wild-type (Berry et al. 2023). As such, it is possible that older *eat-2;mxl-2* animals might not be able to maintain a normal respiratory rate.

**Figure 5.**
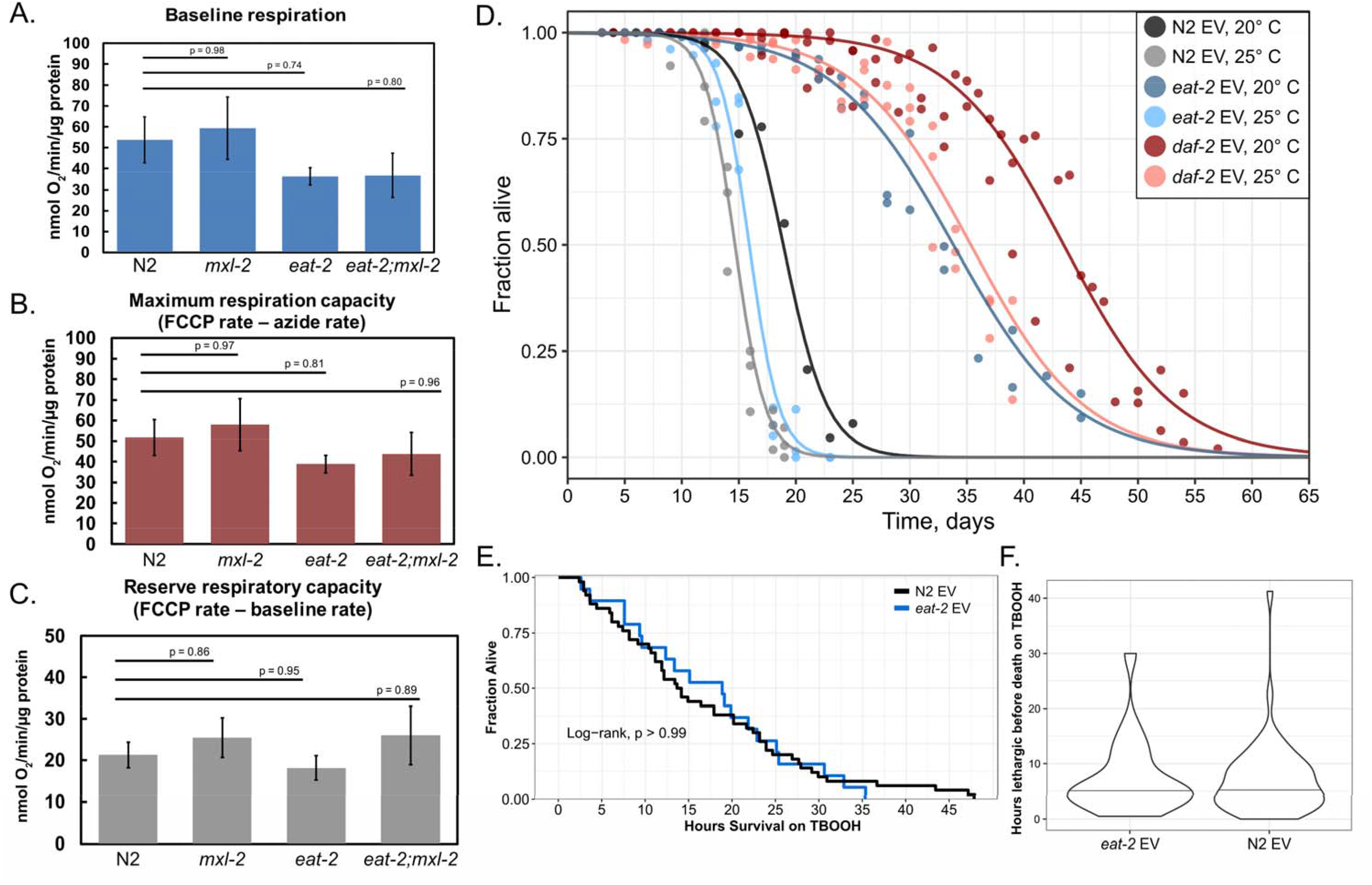
DR in *eat-2(ad465)* animals does not compromise respiratory rate, but also does not confer resistance to oxidative or chronic mild thermal stress. **A-C :** Young adult *eat-2(ad465)* animals do not exhibit reduced respiratory rate, which remains unaltered by loss of *mxl-2* in *eat-2;mxl-2* double mutant animals. Oxygen consumption of whole animals was measured in a Clark-type electrode, and no significant differences were observed between strains at day 2 adult. Oxygen consumption rates are normalized by total protein. Results show mean ± SEM from three biological replicates for each strain, each from a separate synchronized population, and p-values from statistical analysis with one-way ANOVA followed by the Tukey post-hoc test. **(A)** Baseline respiratory rate before treatment with FCCP or sodium azide. **(B, C)** Animals were treated with the oxidative phosphorylation uncoupler FCCP to induce maximal respiratory rate, followed by sodium azide as an inhibitor. The difference between the uncoupled rate and the inhibited rate is the maximum respiratory capacity (B), and the difference between the uncoupled rate and the baseline rate is the reserve capacity under typical conditions (C). **(D)** *eat-2(ad465)* animals are not long-lived under chronic mild heat stress, unlike with reduced insulin signaling in *daf-2*. Replica-set lifespan experiments for N2, *eat-2(ad465)*, and *daf-2(e1370)* mutant animals kept at 20 and 25 °C post-development. Points represent observations of independent sub-populations of animals, fit with a logistic function to obtain a survival curve (Cornwell et al. 2018, 2022). *daf-2* mutant animals have been well-characterized as long-lived and stress-resistant. In contrast, *eat-2* DR animals are long-lived at 20 °C- a typical laboratory culture condition-but not at 25 °C, which has been found to induce a mild heat-stress response (Lee and Kenyon 2009; Gouvea et al. 2015). The addition of chronic mild heat stress completely suppresses *eat-2* lifespan, similar to a previously reported finding (Henderson et al. 2006). **E-F:** Young adult (day 2) *eat-2(ad465)* animals are not more resistant to oxidative stress than wild-type, using treatment with tert-butyl hydroperoxide (TBOOH). Data represents two biological replicate plates per condition, wild-type N2 n = 50 and *eat-2* n = 19. **(E)** No significant difference was found in median survival duration in DR animals, indicating *eat-2* does not provide increased resistance to oxidative stress in young adults. Data were analyzed non-parametrically with Kaplan-Meier, followed by the log-rank test in R. **(F)** Using automated longitudinal imaging, we were able to discriminate between the time when animals cease broad locomotion, and when they eventually die. Similar to survival, we also find no difference in the duration of lethargy proceeding death for wild-type (N2) and *eat-2* animals under oxidative stress. Wilcoxon rank-sum test p-value = 0.8193.

### DR in *eat-2* increases lifespan without a concomitant increase in stress resistance

We next sought to determine whether *mxl-2* was required for enhanced stress resistance of *eat-2* mutant animals. Stress resistance broadly declines during normal aging (Labbadia and Morimoto 2015; Dues et al. 2016; Vos et al. 2016; Son et al. 2019); *eat-2* mutant animals have previously been shown to have increased resistance to proteotoxic stress in aggregation-prone models (Steinkraus et al. 2008; Matai et al. 2019). We chose to assess how a chronic state of DR would impact mild but persistent proteotoxic stress; wild-type animals maintained at 25 °C are slightly short-lived, have accelerated proteostatic decline, but lack the severe fecundity deficits of animals maintained at higher temperatures (Byerly et al. 1976; Gouvea et al. 2015). We previously showed that maintaining wild-type *C. elegans* at 25 °C is sufficient to induce a mild hormetic heat shock (Das et al. 2017); we reasoned if chronic DR extended longevity through improved stress resistance, then *eat-2* mutant animals should have a similar increase in lifespan at 25 °C. To our surprise, we found that keeping *eat-*2 animals at 25 °C instead of 20 °C is sufficient to completely suppress DR lifespan (Figure 5D, Table S4, suggesting chronic DR does not extend longevity via improved resiliency to heat stress.

Dietary restriction reduces oxidative stress in rodent models, and even biomarkers of oxidative stress in humans (Sohal and Weindruch 1996; Il’yasova et al. 2018). To test whether *eat-2* mutant animals had increased resistance to acute oxidative stress, we treated animals with tert-Buytl-hydroperoxide (TBOOH) and assayed survival. To our surprise, we found no significant difference in median survival between wild-type and *eat-2* animals (Figure 5E). To determine whether a more subtle improvement was occurring, we ascertained the time at which animals cease large translational body movements and enter a lethargic state for a period in advance of death. *eat-2* animals under oxidative stress were similarly not more active compared to wild-type (Figure 5F). Thus, it appears that *eat-2* animals do not exhibit improved oxidative stress resistance (OSR) by day 2 adulthood.

Autophagy is critical for maintenance of organismal health, both under stress and normal conditions, and has also been implicated in DR in *C. elegans*, specifically in the intestine (Hansen et al. 2008; Lapierre et al. 2015; Gelino et al. 2016). We were thus surprised to find a lack of transcriptional evidence for autophagy induction in our *eat-2* animals (Figure S7). Even though *eat-2* expressed genes associated with autophagy machinery at wild-type levels, when combined with *mxl-2* mutation, we found significant down-regulation of two genes associated with autophagy: *lgg-1* (orthologous to LC3/Atg8), an important component of autophagosomes involved in cargo recruitment, and *unc-51* (orthologous to ULK1/2) which initiates autophagosome formation; both *lgg-1* and *unc-51* are necessary for long lifespan in a different *eat-2* mutant (Tóth et al. 2008; Meléndez and Levine 2009; Gelino et al. 2016; Lin and Hurley 2016). Interestingly, *daf-16* RNAi in *eat-2;mxl-2* animals ameliorated the down-regulation of *lgg-1*, while *pha-4* RNAi returned *unc-51* to WT-like levels. On the other hand, *daf-16* knockdown in *eat-2;mxl-2* yielded additional downregulation of *let-363* (TOR), *vps-15*, and *atg-9*; while the latter two genes are part of the autophagy machinery they do not appear to influence longevity, whereas *let-363*-like its mammalian homolog TOR-is a key node in the switch between anabolism and catabolism and negatively regulates autophagy (Laplante and Sabatini 2012; Antikainen et al. 2017; Blackwell et al. 2019). Thus, *eat-2* mutant animals do not increase expression of autophagy genes by day two of adulthood, but *mxl-2* is still required for maintenance of critical autophagy gene expression specifically in DR. It remains unclear if *daf-16*(RNAi) has functional consequences on autophagy activity through the downregulation of TOR. While others have shown requirement of autophagy genes for *eat-2* lifespan and an increase of LGG-1 puncta and autophagosome turnover in a different allele of *eat-*2, *ad1116* (Hansen et al. 2008; Gelino et al. 2016), we posit that while basal autophagy activity is maintained in *eat-2*, by early adulthood most available metabolic resources have been mobilized and energy metabolism has been optimized such that further induction of autophagy is no longer a part of a transcriptional adaptive response.

Overall, our data aligns with previous work indicating that *eat-2(ad465)* animals are not as broadly stress-resistant as other long-lived mutants (Kaeberlein et al. 2006; Dues et al. 2019). Intriguingly, *eat-2(ad1116)* and *eat-2(ad453)* animals exhibit an even slower pumping and feeding rate than *eat-2(ad465)* (Lakowski and Hekimi 1998; Boyd et al. 2007) and enhanced resistance to thermal and oxidative stress (Hansen et al. 2007; Shpigel et al. 2019; Soo et al. 2023), which may parallel experiments that have shown a non-linear relationship between the degree of DR and the resulting aging benefits (Mair et al. 2009). In our hands, stress resistance of *eat-2(ad465)* animals is similar to wild-type animals, indicating that lifespan and healthspan benefits from chronic and constitutive restricted food availability in this model do not arise from raising general barriers against stress.

### Loss of MXL-2 yields synthetic dysregulation of gene expression, inverting the expression of hundreds of genes normally repressed in DR

Strikingly, the *eat-2;mxl-2* double mutant animals yielded two distinct gene expression patterns. First, we observed a synthetic gene expression phenotype: a number of genes expressed at wild-type levels in either *eat-2* or *mxl-2* single mutant animals, were up-regulated in *eat-2;mxl-2* (Figure 2D, Table S5). 1207 and 1347 genes were synthetically up- or down-regulated in *eat-2;mxl-2*, respectively. Second, we observed a pattern of inversion in expression within *eat-2;mxl-2* animals: 507 genes normally repressed in *eat-2* become significantly upregulated in *eat-2;mxl-2* (Figure 6A, Table S5). In contrast, inactivation of *pha-4* in DR animals results in the majority of genes that were previously differentially down-regulated in *eat-2* reverting to basal wild-type-like levels, and without synthetic changes in expression (Figure 6B). A schematic illustration of the synthetic and inverted genetic interactions is illustrated in Figure 6E.

**Figure 6.**
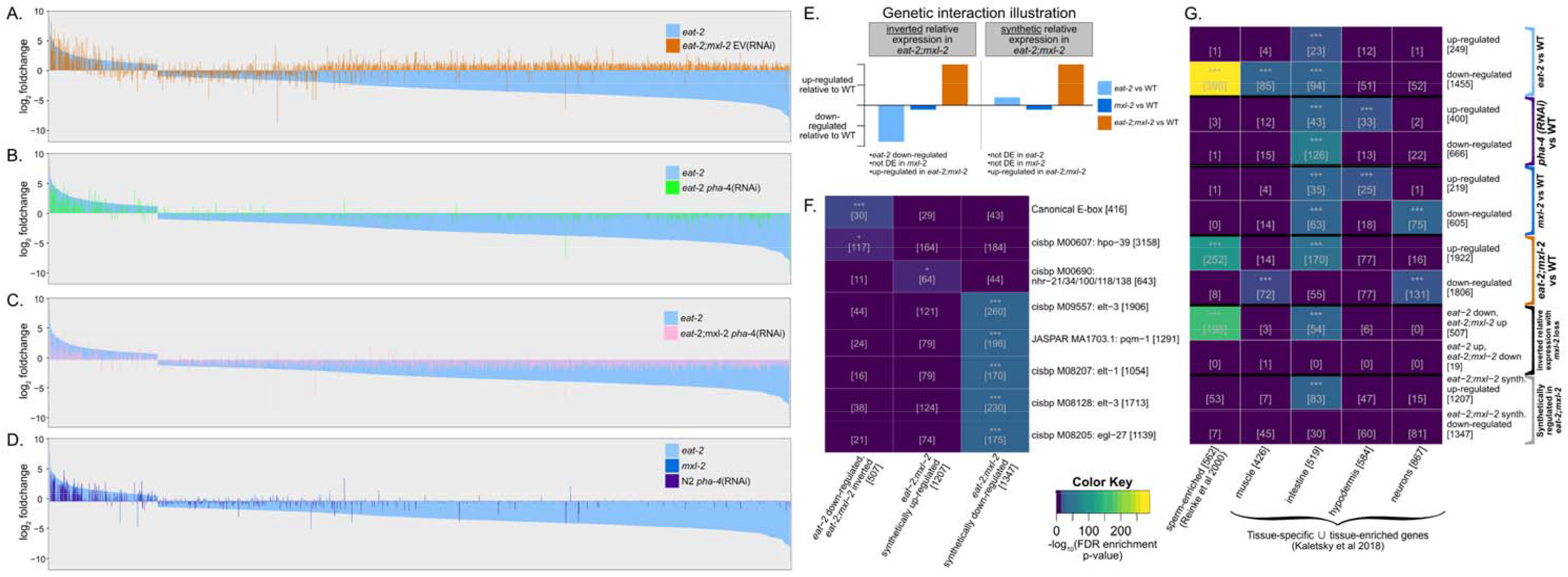
Loss of *mxl-2* in the context of DR results in significant upregulation of 507 genes normally downregulated in *eat-2*. DR, as well as *pha-4* and *mxl-2* loss alone, modulate expression of intestinally-enriched genes, but *eat-2* animals also show down-regulation of genes enriched in sperm and muscle. Sperm, but not muscle genes, are synthetically up-regulated in *eat-2;mxl-2* double mutant animals. **A-D**: *All eat-2* DE signature genes ordered by foldchange in *eat-2*, represented by the light blue bars; the jump between up- and down-regulated genes is due to the foldchange cutoff applied to determine the DE gene signature. Foldchange for the same gene in the indicated comparison versus wild-type (N2 EV) is overlaid: **(A)** *eat-2;mxl-2*, **(B)** *eat-2 pha-4(RNAi)*, **(C)** *eat-2;mxl-2 pha-4(RNAi)*, and **(D)** *mxl-2* or *pha-4* loss in otherwise wild-type animals. Surprisingly, we find that 507 genes significantly down-regulated in eat-2 are significantly up-regulated in *eat-2;mxl-2* double mutant animals, representing almost 35% of all genes downregulated in DR (A). The same pattern is not observed with DR upregulated genes, or with pha-4 RNAi *in eat-2* animals (A, B). (C.) Inactivation of *pha-4* via RNAi in *eat-2;mxl-2* mutants animals suppresses the “inversion” of relative expression observed in *eat-*2;*mxl-2* down-regulated genes, instead broadly resembling *eat-2 pha-4(RNAi)*. **(E.)** Schematic illustration of categories of significant gene expression changes we found in *eat-2;mxl-2* animals that were not observed in *eat-2* or *mxl-2* single-mutant animals alone. Cases where a gene was significantly different from WT in *eat-2* animals, and significantly differentially expressed in the opposite direction relative to WT in *eat-2;mxl-2* animals are hereafter referred to as *eat-2* genes with inverted relative expression in *eat-2;mxl-2* (left side). We further find a large class of genes with significant expression changes in *eat-2;mxl-2* compared to WT, but remain at WT-like levels in *eat-2* and *mxl-2*, and these are hereafter referred to as having synthetic relative expression in *eat-2;mxl-2* (right side). **(F.)** Top hits for TF motif-based predicted binding enrichment of the sets of genes with inverted relative expression between *eat-2* and *eat-2;mxl-2* or genes synthetically differentially expressed in *eat-2;mxl-2.* Genes which are down-regulated in *eat-2* but significantly up-regulated in *eat-2;mxl-2* are enriched for the canonical CACGTG E-box E-box sites in their promoter regions, and for motif matches to the bZIP TF *hpo-39*. In contrast, genes synthetically upregulated in *eat-2;mxl-2* have enrichment for promoter matches to the motif of nuclear hormone receptor family paralogs *nhr-21*, *34*, *100*, *118*, and *138*, and the synthetically down-regulated genes in *eat-2;mxl-2* exhibit substantial enrichment for predicted promoter binding of GATA factors *elt-1*, *elt-3*, *egl-27*, and *pqm-1*. Note that two separate experiments have identified motifs for ELT-3, they are both shown. **(G.)** Tissue enrichment for significantly differentially expressed in genes in *eat-2* EV, *mxl-2* EV, N2 *pha-4(RNAi)*, *eat-2;mxl-2* EV, all compared to wild-type (N2 EV). Also including tissue enrichment for the genes with inverted relative expression between DR (*eat-2*) and *eat-2;mxl-2* double mutant animals, as highlighted in (A), and the broader set of synthetic gene expression changes in eat-2;mxl-2 but not in single-mutants. We found that these synthetically regulated genes– and particularly those that are normally down-regulated in DR– are enriched in the intestine and in sperm. Somatic tissue gene sets are based on the union of tissue-specific and tissue-enriched gene lists from cell type specific bulk RNA-Seq in (Kaletsky et al. 2018). Sperm-specific genes are from (Reinke et al 2000). Numbers in brackets indicate the size of the gene set, or the number of genes in the intersection. The color key is the same for panels **F** and **G,** and stars also indicate the hypergeometric test adjusted p-value significance level in both cases: ‘***’ p < 0.001 ‘**’ 0.01 ‘*’ 0.05 ‘.’ 0.1.

### Opposing Myc-family members, Nuclear Hormone Receptors, and GATA TFs putatively regulate changes in gene expression when MLX-2 is unable to respond to chronic DR

We reasoned the aforementioned inverted and synthetic changes in gene expression may be revealing novel transcriptional responses that are germane to understanding how biological systems maintain homeostasis under conditions of metabolic stress when one regulatory component is impaired. To identify putative upstream regulators and tissue-associated biases in expression for the inverted and synthetic up or down gene sets, we analyzed each set for over-representation of TF binding sites in promoters and tissue specificity. As only a fraction of the known *C. elegans* transcription factors have been profiled via ChIP-Seq, protein-binding microarrays, or newer approaches such as CUT&TAG to map context-specific binding sites, we were able to perform *in silico* binding predictions based on motif matching for more than one-third of *C. elegans* TFs (Fuxman Bass et al. 2016). Concurrently, we analyzed in more detail the presence of E-box sequences within the *C. elegans* genome, as the Myc-family of TFs bind to these sequences. In *C. elegans* the MML-1::MXL-2 and MDL-1::MXL-1 complexes bind E-box motifs (CACGTG or CANNTG), (Pickett et al. 2007; Grove et al. 2009; Gerstein et al. 2010). Additionally, we included manually-defined motifs for the “canonical” CACGTG E-box, a degenerate E-box motif defined based on MYC::MAX heterodimer binding (Allevato et al. 2017), and the carbohydrate response element (ChoRE) motif bound by the mammalian homolog of MML-1, ChREBP/MONDOB/MLXIPL (Shih et al. 1995). Overall, we found 695 genes with E-box-like motifs in promoter regions when restricted to matches also found in homologous genes of at least two other nematode species, and 3108 gene matches in *C. elegans* when homology was not considered; about half of these matches were for the canonical CACGTG E-box (Table S6). We are not aware of any previous effort to characterize the presence of ChoRE motifs in the *C. elegans* genome, and were surprised to find just 397 matches, which were not enriched for any pathway in KEGG or Reactome, or Gene Ontology term. However, *mdl-1* and *mxl-3* were among the 387 genes identified. This leaves open the remote possibility that MML-1 regulates expression from genes with hitherto unidentified ChoRE-like motifs in *C. elegans*.

As proof-of-principle, we first analyzed single genetic perturbations and found specificity in predicted motif sites among the promoters of down-regulated genes. Three different E-box motifs were enriched among all down-regulated genes in *eat-2* animals: canonical CACGTG E-boxes, MDL-1 and CRH-2 (a bHLH transcription factor known to bind at degenerate E-box sites) (Figure S8A). As expected, the *daf-16* DNA binding motif was the most significantly enriched with *daf-16(RNAi)* treatment. Neither PHA-4 nor E-box sites were enriched in the down-regulated DE genes of *pha-4(RNAi)* or *mxl-2(0)* animals, which may be due to limited activity of these factors under well-fed conditions (Sheaffer et al. 2008; Johnson et al. 2014). Interestingly, we did not observe enrichment of PHA-4 sites even in the subset of *pha-4*-dependent DR genes, indicating an indirect regulatory role for PHA-4 in *eat-2*, PHA-4 binding at motifs that are distinct from those that have already been characterized, or that PHA-4 acts primarily at earlier developmental stages-this is consistent with the role of PHA-4(FOXA) as a pioneer TF, which keep enhancer nucleosomes accessible in chromatin, thereby allowing other TFs to bind and regulate transcription (Zaret and Carroll 2011; Zaret and Mango 2016). We were encouraged that significant enrichment for TF binding motifs were observed under conditions in which one would expect changes in TF activity.

Across known TF DNA binding motifs, we found distinct significant TF motif enrichment in the three gene sets (i.e., inverted from *eat-2* with *mxl-2* loss, synthetically upregulated or downregulated in *eat-2;mxl-2*). Enrichment for predicted transcription factor motif binding identified significant overlap for the canonical CACGTG E-box in the inverted gene set; tissue-associated enrichment revealed significant expression in sperm and intestine (Figure 6F-G). Transcriptionally activating and repressing heterodimers of bHLH Myc-family TFs are known to compete for binding at E-box elements (Pickett et al. 2007; Conacci-Sorrell et al. 2014), and previous results suggested that the MML-1::MXL-2 and MDL-1::MXL-1 heterodimeric complexes may function as a rheostat in the transcriptional regulation of longevity (Johnson et al. 2014). We posit that loss of *mxl-2*, and therefore MML-1::MXL-2 complex binding, facilitates MDL-1::MXL-1 binding in *eat-2* for the loci yielding the inverted gene expression pattern. Collectively, it is tempting to speculate that the convergence of opposing Myc-family heterodimers at these loci may be key for coupling metabolic changes to reproductive fecundity.

We next examined the synthetically differentially expressed genes (i.e., DE in *eat-2;mxl-* 2, but not either single mutant). In contrast to the inverted gene set, synthetically up-regulated genes failed to have significant enrichment for E-box sequences, suggesting regulation by TFs other than Myc-family members. Significant motif enrichment for synthetically up-regulated genes was limited to a subset of nuclear hormone receptor paralogs-*nhr-21, 34, 100, 118, 138*- and enriched for predicted intestinal expression (Figure 6F-G). Lastly, significant enrichment for GATA binding factors- *elt-1*, *elt-3*, *pqm-1*, and *egl-27*- were found in genes synthetically down-regulated (Figure 6F). Interestingly, *pqm-1* is required for *eat-2* lifespan, and acts antagonistically with DAF-16 for nuclear entry (Tepper et al. 2013; Shpigel et al. 2019). Whether or not these TFs functionally impact longevity under these conditions is unclear; yet a picture begins to emerge. Loss of *pha-4* completely suppresses *eat-2* lifespan (i.e., the lifespan of *pha-4(RNAi)* and *eat-2;pha-4* are the same) (Panowski et al. 2007). In contrast, we find that while loss of *mxl-2* significantly shortens *eat-2* lifespan, suppression is only partial (i.e., lifespan of *mxl-2(0)* is shorter than *eat-2;mxl-2*) (Johnson et al. 2014). Yet loss of either *pha-4* or *mxl-2* is sufficient to suppress the vast majority of the DE signature we observe in *eat-2* mutant animals, but only loss of *mxl-2* produces synthetic changes in gene expression in animals undergoing chronic metabolic stress. It remains to be determined if NHR and GATA transcription factors are activated in an attempt to compensate for the breakdown in appropriate regulation of gene expression in chronic DR without *mxl-2*, or if this represents dysfunction of a transcriptional network in which MML-1 and MXL-2 normally play crucial roles. In future studies it will be interesting to explore these possibilities in greater mechanistic detail.

## Discussion

It has been known for centuries that caloric restriction extends longevity. Luigi Cornaro the Venetian wrote four *Discorsi* between 1550 and 1562; describing how his adoption of a temperate lifestyle and consuming only 350 grams of food daily were the basis for his attaining a long and healthful life (Cornaro 1727, 1905). Since the modern era of genetic discovery, indeed a common theme has emerged: evolutionarily conserved genes and pathways that regulate the progression of longevity and those with the largest impact often act as “watchtowers” of nutrient and energy availability (Lazaro-Pena et al. 2022). Why would there be deep evolutionary conservation causally linking mechanisms of nutrient sensing to organismal longevity? We posit that organisms able to couple physiology to energy resources had a survival advantage when food is scarce by conserving and recycling resources, while delaying energetically-costly physiological processes, such as development and reproduction. We find the transcriptional signature of *eat-2* mutant animals, a form of constitutive and chronic DR, broadly involves the down-regulation of gene expression associated with amino acid and lipid metabolism, and other energetically expensive processes such as collagen production and muscle mass.

Delaying the production of offspring has the benefit of limiting competition for limited resources. In *C. elegans* many long-lived mutant animals, including those with *eat-2* mutations, have slower development, reduced numbers of overall progeny, and an extended period of progeny production (Scharf et al. 2021). We find downregulation of genes that encode functions related to sperm production in *eat-2* mutant animals were dependent upon *mxl-2*; in the absence of *mxl-2* down-regulation of gene expression fails to occur, brood size from self-progeny is reduced, and non-viable embryos are produced. Thus, our work reveals that MXL-2 plays a key role in mediating a transcriptional adaptive response that links energy availability to reproductive fitness. *C. elegans* hermaphrodites are limited to producing only a limited number of sperm: approximately 300, but mated animals can produce nearly 1400 progeny (Singson 2001). In response to limited food availability decreased expression of genes required to generate sperm could provide a simple but elegant method maximize reproductive fitness by limiting total number of sperm. Importantly, refined genetic analysis revealed that alterations in reproduction/development are separable from longevity. For example, early discoveries in *C. elegans* aging research using temperature-sensitive alleles in the insulin/IGF1 pathway (IIS) revealed that dauer formation and extended longevity were genetically separable (Kenyon et al. 1993), which implies strategies to improve healthy aging may not require a cost in developmental or reproductive fitness.

It has been suggested *eat-2* animals are a model of dietary restriction when raised on *E. coli* due to intestinal bacteria colonization resulting from insufficient pharyngeal grinder function, in turn leading to upregulation of innate immune responses and food-avoidance behavior (Kumar et al. 2019). While we see a handful of innate-immune-related genes upregulated, we find that set is not specific to *eat-2*. Thus, gene expression changes in innate immunity are unlikely to be the result of intestinal colonization, but rather a low-level compensatory response to perturbations that affect the intestine. We have previously established that *mxl-2* animals do not have pharyngeal pumping defects (Johnson et al. 2014), and are indistinguishable from wild-type with respect to adult size and development rate. It seems unlikely that infection response is regulating gene expression in DR differently from other contexts. However, Myc-factor function has been demonstrated with response to the microsporidian intestinal pathogen *N. parisii*, wherein MXL-2 appeared to be functioning separately and non-canonically from MML-1: loss of *mdl-1, mxl-1,* or *mxl-2* alleviated infection severity, while loss of *mml-1* increased it (Botts et al. 2016). Whether these responses connect to DR signals by relaying intestinal stress remains unknown.

Different methods of generating DR in *C. elegans*, either through alteration in food availability, type or amount of bacteria, specific nutrient restriction, timing of restriction, or genetic perturbation have vastly different genetic requirements for DR benefits (Greer and Brunet 2009; Denzel et al. 2018). In comparison to non-genetic DR models in *C. elegans*, *eat-2* differs substantially in that DR in *eat-2* is constitutive and chronic, even when multiple generations are raised on adequate food. Genetic perturbation of the links between nutrient sensing and adaptive responses that extend longevity have a high degree of specificity to the nutrient, tissue of action, and cell-type (Templeman and Murphy 2018; Miller et al. 2020). For instance, TOR responds to levels of amino acids and carbohydrates to switch between cellular growth and maintenance and longevity (Jia et al. 2004; Hansen et al. 2008; Kapahi et al. 2010; Seo et al. 2013; Lapierre et al. 2015); AMPK is a conserved energy sensor of increased levels of AMP and ADP, and induces autophagy, the oxidative stress response (OSR), and extends longevity (Apfeld et al. 2005; Greer et al. 2007; Greer and Brunet 2009; Salminen and Kaarniranta 2012); Sirtuins, a family of (NAD+)-dependent deacetylases, sense levels of NAD+ to regulate longevity (Jedrusik-Bode et al. 2013; Jedrusik-Bode 2014; Verdin 2015); and genetic perturbations that decrease IIS delay aging from *C. elegans* to humans (Barbieri et al. 2003; Taguchi and White 2008). The aforementioned metabolic-longevity signals converge on a limited number of transcription factors, including *skn-1* (ortholog of nuclear factor erythroid 2-related factor 2, Nrf2), *daf-16* (FOXO), *pha-4* (FOXA), *mml-1, mxl-2, hsf-1* (heat shock transcription factor), *hif-1* (HIF1), and *hlh-30* (TFEB) (Lin et al. 1997; Sheaffer et al. 2008; Tullet et al. 2008; Zhang et al. 2009; Lapierre et al. 2013; Johnson et al. 2014; Nakamura et al. 2016; Lazaro-Pena et al. 2022) and direct transcriptional regulators, such as *hpk-1* (homeodomain-interacting protein kinase) (Das et al. 2017; Lazaro-Pena et al. 2023). Collectively, a growing number of *C. elegans* studies have begun to unravel the complex integrated networks that maintain organismal homeostasis from an extensive array of diverse extrinsic and intrinsic signals that converge on distinct but overlapping adaptive transcriptional responses (Greer and Brunet 2009; Denzel et al. 2019). All the aforementioned signal transduction pathways act acutely via monitoring metabolic state; we find that *eat-2* mutant animals regulate a distinct transcriptional program through MXL-2 to maintain homeostasis under conditions of long-term metabolic stress. It would be interesting in the future to create a conditional allele of *eat-*2, for example via CRISPR-tagging endogenous *eat-2* with the Auxin-Inducible Degradation (AID) system (Zhang et al. 2015), thereby creating a generalizable genetic model of dietary restriction to further elucidate how these myriad of metabolic watchtowers are integrated within multicellular organisms.

The *C. elegans* Myc-network is a key integration point for multiple longevity signals, including decreased IIS, decreased pharyngeal pumping and feeding of *eat-2* mutants, TORC1 inhibition, *glp-1* mutant germline-less animals, and HLH-30 (TFEB). All activate the MML-1::MXL-2 complex by promoting nuclear accumulation of MML-1 (Johnson et al. 2014; Nakamura et al. 2016; Shioda et al. 2023), consistent with known modes of regulation of mammalian MLXIP/MLXIPL and MLX (Billin and Ayer 2006; Sloan and Ayer 2010; Diolaiti et al. 2015). We previously discovered that MML-1::MXL-2 complex function is necessary for the increased lifespan of not only *daf-2* and *eat-2* mutant animals, but also oxidative stress resistance and heat shock survival (Johnson et al. 2014). The MML-1::MXL-2 complex is also essential for the increased longevity conferred by TORC1 inhibition (Nakamura et al. 2016; Lazaro-Pena et al. 2023). MML-1::MXL-2 is also necessary for *hpk-1*, an essential component of TORC1-mediated longevity, to increase lifespan, induce autophagy and maintain proteostasis during aging (Das et al. 2017; Lazaro-Pena et al. 2023). Previous work characterizing stress resistance in *eat-2* found that DR buffers against proteotoxic stress, specifically the accumulation of toxic aggregates (Steinkraus et al. 2008; Matai et al. 2019; Shpigel et al. 2019); however the *eat-2* mutation does not protect against chronic mild thermal stress ((Kaeberlein et al. 2006) and this study). We also find a lack of broad stress resistance in *eat-2* mutant animals, nor transcriptional changes associated with stress resistance, despite a significantly longer lifespan and healthspan. This supports the notion that the Myc-family of transcription factors not only integrate longevity signals (Johnson et al. 2014; Nakamura et al. 2016), respond to acute forms of metabolic stress, but coordinate unique signal-specific adaptive responses and play a key role in transcriptional remodeling to extended periods of dietary deprivation.

## Author Contributions

A.B.C designed and conducted genomic analyses, performed survival experiments, and analyzed all experimental data. M.T. prepared animal populations and isolated RNA for RNA-Sequencing. Y.Z. designed and conducted brood size experiments. J.T. advised genomic analyses and assisted with development of the manuscript. A.V.S. oversaw the project, contributed to analysis and interpretation of results, and obtained funding. A.B.C. and A.V.S. wrote the manuscript.

## Acknowledgements

Some strains were provided by the CGC, which is funded by NIH Office of Research Infrastructure Programs (P40 OD010440), and by the National BioResource Project, Japan. We would like to acknowledge the UR Center for Integrated Research Computing (CIRC), the University of Rochester Genomics Research Center (GRC) for their resources and support. We would also like to thank members of the Biomedical Genetics department and the Western New York Worm Group for helpful discussions and feedback, particularly Dr. Doug Portman and Dr. Keith Nehrke. The oxygen consumption experiments were performed in conjunction with Brandon Berry of Dr. Andrew Wojtovich’s laboratory.

## Data Availability

The RNA-Seq dataset is available from the Gene Expression Omnibus (GEO) under accession GSE240821.

---

**Supplementary Figure S1.**
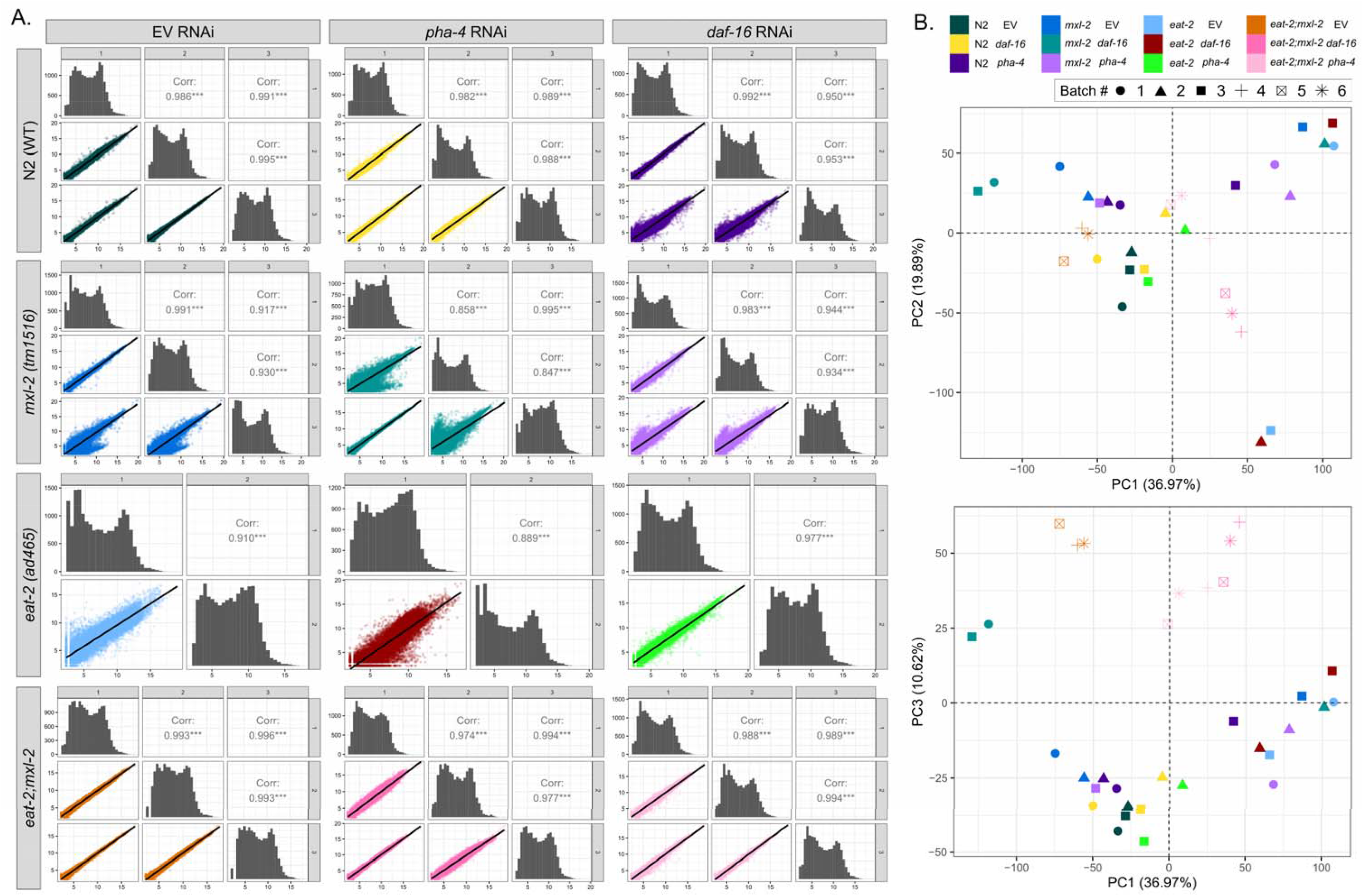
RNA-Seq dataset intra-group replicate sample correlations are strong, and variability is not associated with known batches. (A.) Pearson correlation for replicate samples within sample groups is generally strong, and not lower than 0.847 in any case. Sample groups are organized by strain as indicated on the left side, and RNAi condition as indicated on the top. All sample groups have two or three replicates. Histograms indicate the density of the points in the correlation dot plots. Correlation values are based on Pearson correlation, and all correlations were significant, as indicated by the stars. All 17,907 genes considered as expressed in any sample in the dataset are included. (B.) Due to the size of the experiment, animal samples were prepared in six different batches, as indicated by the shape of the points in the PCA scatterplots. Across the first three principal components, variability between samples is not associated with these batches, and as such we do not find evidence of batch effects. Note that while animals were prepared in separate batches, all samples were sequenced in the same run.

**Supplementary Figure S2.**
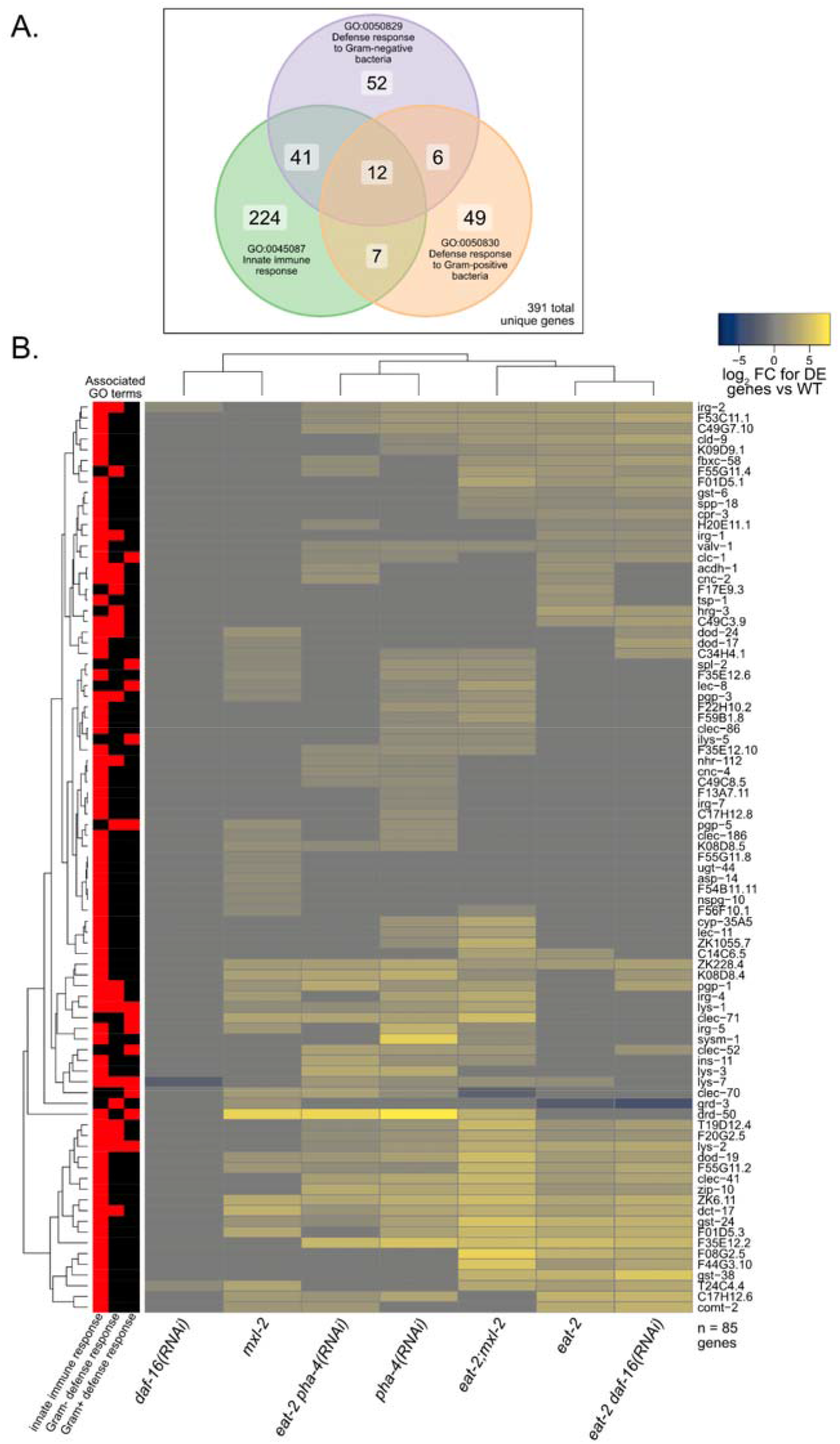
A subset of innate immune response-associated genes are upregulated in *eat-2*, but are not specific to the DR context. As we observed enrichment of innate immune response Gene Ontology (GO) terms in *eat-2* animals (Figure 2C), we further investigated the relative expression for innate immune response-associated genes in our other sample groups. We find the upregulation of innate immune genes is not specific to the context of *eat-2* and also occurs in *mxl-2* mutants and with *pha-4* RNAi in otherwise basal conditions. **(A)** Venn diagram indicating the number of *C. elegans* genes associated to GO terms for innate immune response, derived from annotation in WormBase WS282. 391 genes are represented in the union of the three innate immune response terms. **(B)** The union of the innate immune response associated genes in (A) was intersected with any of the genes we found significantly up-regulated in *eat-2*, *mxl-2*, or N2 *pha-4* RNAi, yielding 85 genes. The heatmap indicates the foldchanges for the indicated sample group compared to wild-type (N2) for these genes. Red blocks on the left bar indicate the GO terms each gene is associated with. We identified a set of immune response related genes that are similarly upregulated in *eat-2*, *mxl-2*, and N2 *pha-4* RNAi, which may indicate induction of a stress response that is not specific to DR. We further see that the upregulation of some of these genes in *eat-*2 is not affected by loss of *daf-16*, *mxl-2*, or *pha-4*, but that they are not induced by *daf-16* RNAi alone under basal conditions.

**Supplementary Figure S3.**
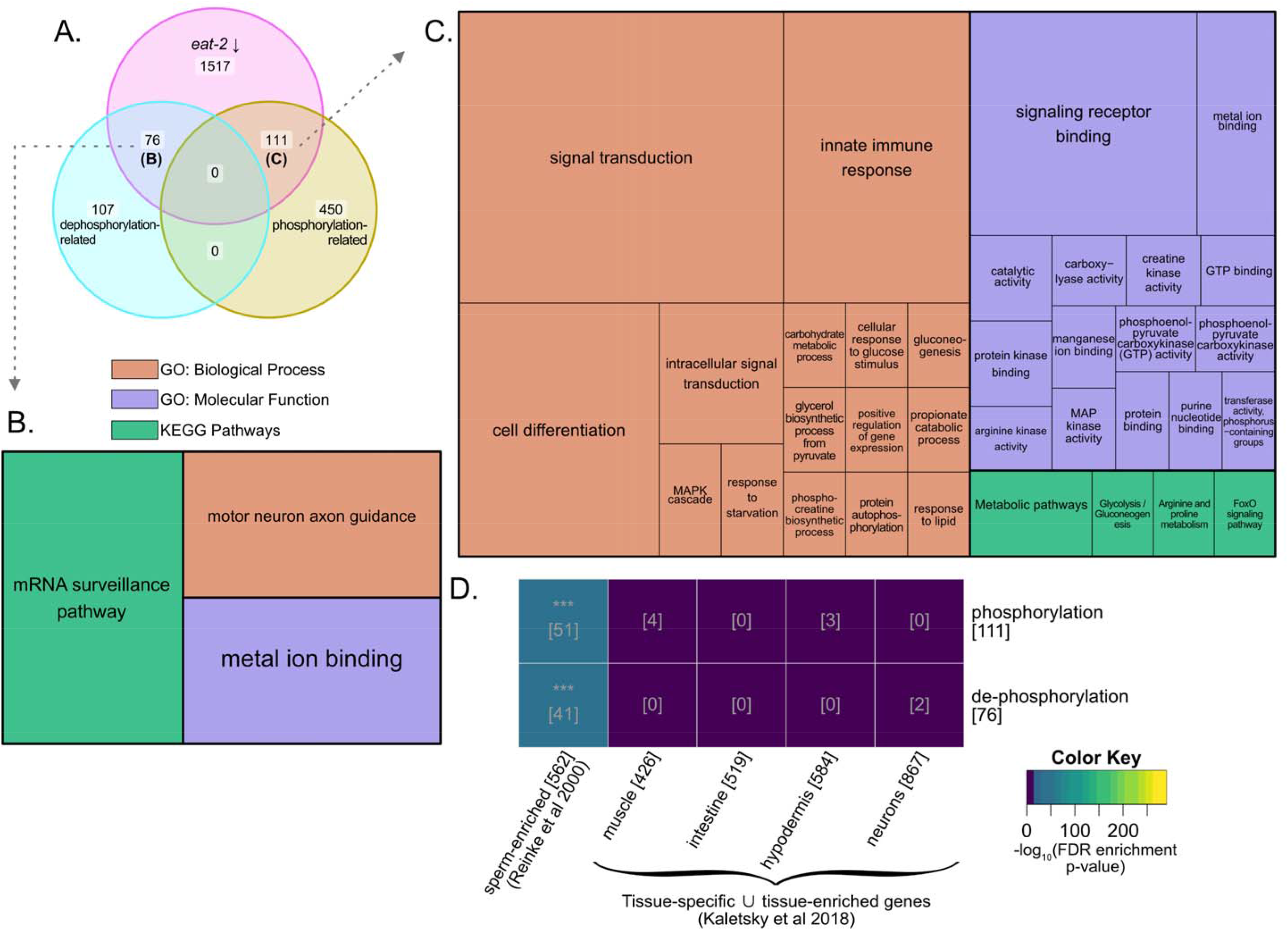
Genes with demonstrated or predicted functions in phosphorylation or de-phosphorylation were significantly enriched among transcripts downregulated in *eat-2*. **(A.)** Based on Gene Ontology term associations from WormBase (WS282) for broad phosphorylation cycle-associated functions, we found 76 de-phosphorylation genes and 111 phosphorylation genes significantly downregulated in *eat-2*. To get a better sense of where these genes may function, or if they participate in signaling cascades, we looked at the other functions and pathways that these genes have been identified in, without considering statistical analysis of the overlaps therein. After removing the broad phosphorylation cycle terms that were used to initially select these genes, and further showing only terms for which more than two genes were associated, we represented the relative frequency of the functional terms as a treemap each for the de-phosphorylation genes **(B.)** and phosphorylation genes **(C.)** downregulated in *eat-2*. In this representation, the size of the box represents the number of genes associated with the term, relative to all the associations for the group. The substantially larger number of functions shown for the phosphorylation genes may be attributable to a larger body of research results on kinases. **(D.)** Tissue enrichment for the phosphorylation and de-phosphorylation genes down-regulated in *eat-2*. Somatic tissue gene sets are based on the union of tissue-specific and tissue-enriched gene lists from cell type specific bulk RNA-Seq in (Kaletsky et al. 2018). Sperm-specific genes are from (Reinke et al. 2000). This shows that both classes of genes are significantly enriched in sperm, rather than being distinctly regulated in different somatic tissues. Numbers in brackets indicate the size of the gene set, or the number of genes in the intersection. Hypergeometric test adjusted p-value significance level: ‘***’ p < 0.001 ‘**’ 0.01 ‘*’ 0.05 ‘.’ 0.1.

**Supplementary Figure S4.**
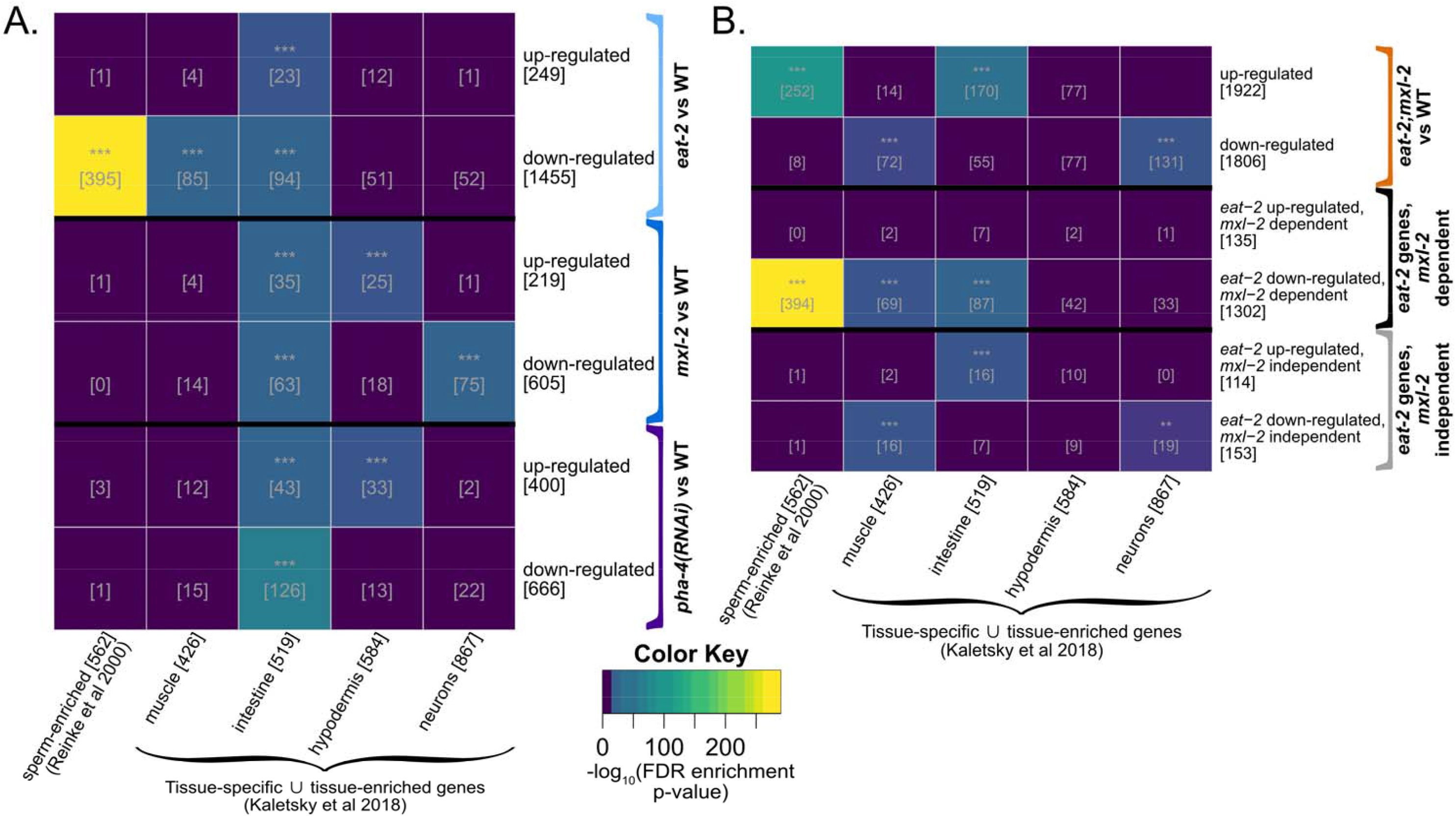
DR conditions, as well as *mxl-2* and *pha-4* loss, modulate intestine-associated gene expression, but *eat-2* animals show down-regulation of sperm and muscle genes, dependent on *mxl-2*. **(A.)** Tissue enrichment for significantly differentially expressed in genes in *eat-2*, *mxl-2*, and *pha-4 RNAi* all compared to wild-type (N2 EV). While regulation of intestine-associated genes is not specific to *eat-2*, we found strong enrichment of muscle- and especially sperm-associated genes in *eat-2* but not loss of *mxl-2* or *pha-4*. **(B.)** We then looked at the set of *eat-2* genes that were dependent on or independent of *mxl-2* and found that 394 of 395 sperm-associated genes downregulated in *eat-2* were dependent on *mxl-2* to maintain reduced expression in DR. Rather than returning to WT-like-levels, a substantial fraction of sperm-related genes are significantly upregulated in *eat-2;mxl-2* animals. Somatic tissue gene sets are based on the union of tissue-specific and tissue-enriched gene lists from cell-type-specific bulk RNA-Seq from (Kaletsky et al 2018). Sperm-specific genes are from (Reinke et al 2000). Numbers in brackets indicate the size of the gene set, or the number of genes in the intersection. Adjusted p-value significance level: ‘***’ p < 0.001 ‘**’ 0.01 ‘*’ 0.05 ‘.’ 0.1.

**Supplementary Figure S5.**
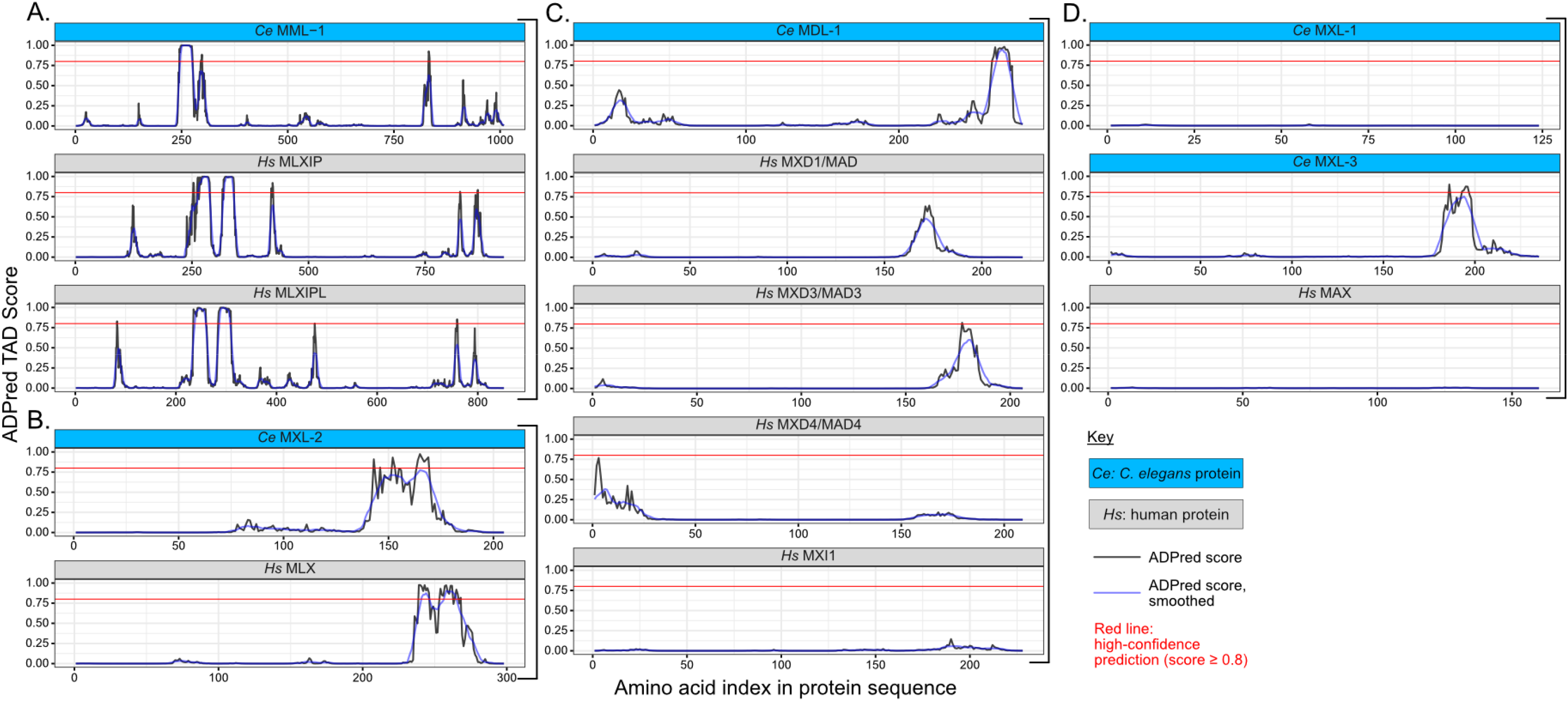
Trans-activation domain (TAD) prediction yields putative high-confidence TAD on the amino-terminal side of MML-1 and the carboxy-terminal side of MDL-1. The presence of trans-activation domains (TADs) in the *C. elegans* Myc-family TFs and their human homologs was predicted with ADpred, a neural-network based model trained on TAD sequences from large-scale screens in yeast (Erijman et al 2020). **(A.)** ADpred analysis identified a high-confidence TAD from 245-276 on the amino terminal side of MML-1, which is between the “Mondo-conserved” domains IV and V (Ceballos, Esse, Grishok 2021) and parallels similar predicted TADs in the human homologs MLXIP and MLXIPL. Note that the MML-1 B isoform differs on the C-terminal end and does not affect the predicted TAD (not shown). **(B.)** We also found potential TAD regions of lower confidence on the C-terminal side of MXL-2, which also correspond to a region of similar confidence on the C-terminal side of human homolog MLX. **(C.)** While *C. elegans* MDL-1 has been thought function primarily as a transcriptional repressor, we found a high confidence TAD predicted on the C-terminal end of MDL-1 as well, indicating that MDL-1 may also have roles in transcriptional activation in some contexts, unlike its human counterparts. **(D.)** Similar to mammalian MAX, we find no evidence of TADs in MXL-1, but do observe a lower confidence predicted TAD on the C-terminal side of MXL-3. The red horizontal line indicates an ADpred score of 0.8, which was suggested as a threshold of high-confidence TAD prediction in (Erijman et al 2020). Protein sequences were from WormBase for *C. elegans* and UniProt for human-MLXIP (Q9HAP2), MXLIPL (Q9NP71), MLX (Q9UH92), MXI1 (P50539), MXD1/MAD (Q05195), MXD3/MAD3 (Q9BW11), MXD4/MAD4 (Q14582), MAX (P61244).

**Supplementary Figure S6.**
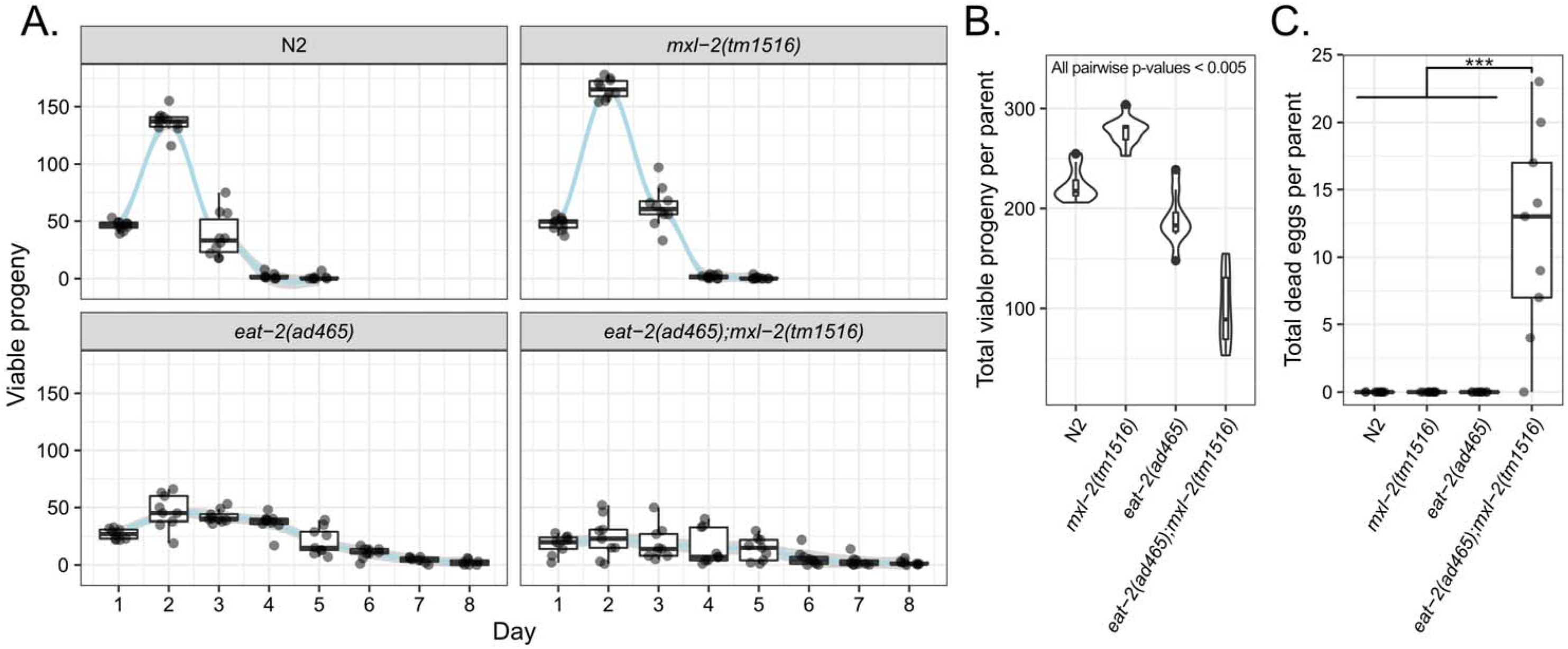
Additional trial for brood size assay. Additional trial data for brood size assay, performed similarly to the representative trial in Figure 5. In this trial as well, we found loss of *mxl-2* in DR but not ad libitum animals compromises fecundity and embryo viability. Brood size assays were performed for N2, mxl-2, eat-2, and eat-2;mxl-2 animals. For each strain, animals were synchronized and parent hermaphrodites were picked to their own plate at L4. Starting from 24 hours after L4, parents were picked to a new plate every 24 hours until no further eggs were observed. Larval animals were counted after 48 hours, allowing enough time for all viable eggs to hatch. Eggs remaining unhatched after this time were considered dead and recorded. Dietarily restricted animals are reproductive through a longer period of their life (”reproductive-span”), as previously reported (Crawford et al 2007, Luo et al 2009), which we find MXL-2 to not be required for **(A)**. However, the smaller brood size typical of eat-2 animals was further reduced in eat-2;mxl-2 double mutants **(B)**. Furthermore, dead eggs were observed for double mutants, but not the other strains **(C)**. See Table S3 for statistical analysis.

**Supplementary Figure S7.**
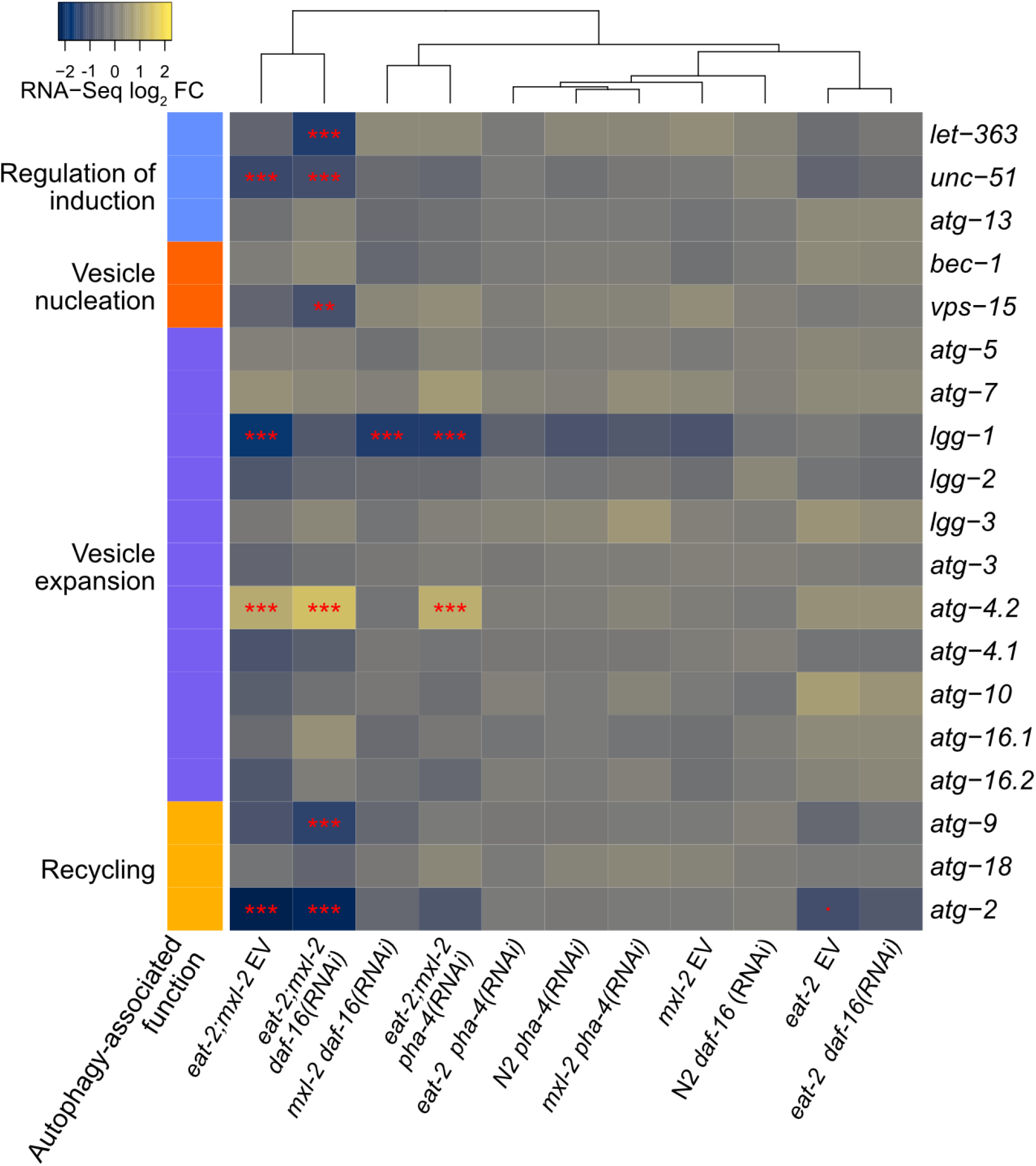
Differential expression results across all comparisons for key autophagy-associated genes. Heatmap indicates the log_2_ foldchanges for the indicated sample group compared to wild-type (N2) for key genes involved in autophagy, with their canonical role indicated by the left colored side-bar. Autophagy-related gene expression was not found to be up-regulated in *eat-2* animals, but was dysregulated by loss of *mxl-2* in the *eat-2* background. Significantly differentially expressed genes are indicated by red stars, adjusted p-value significance level: ‘***’ p < 0.001 ‘**’ 0.01 ‘*’ 0.05 ‘.’ 0.1.

**Supplementary Figure S8.**
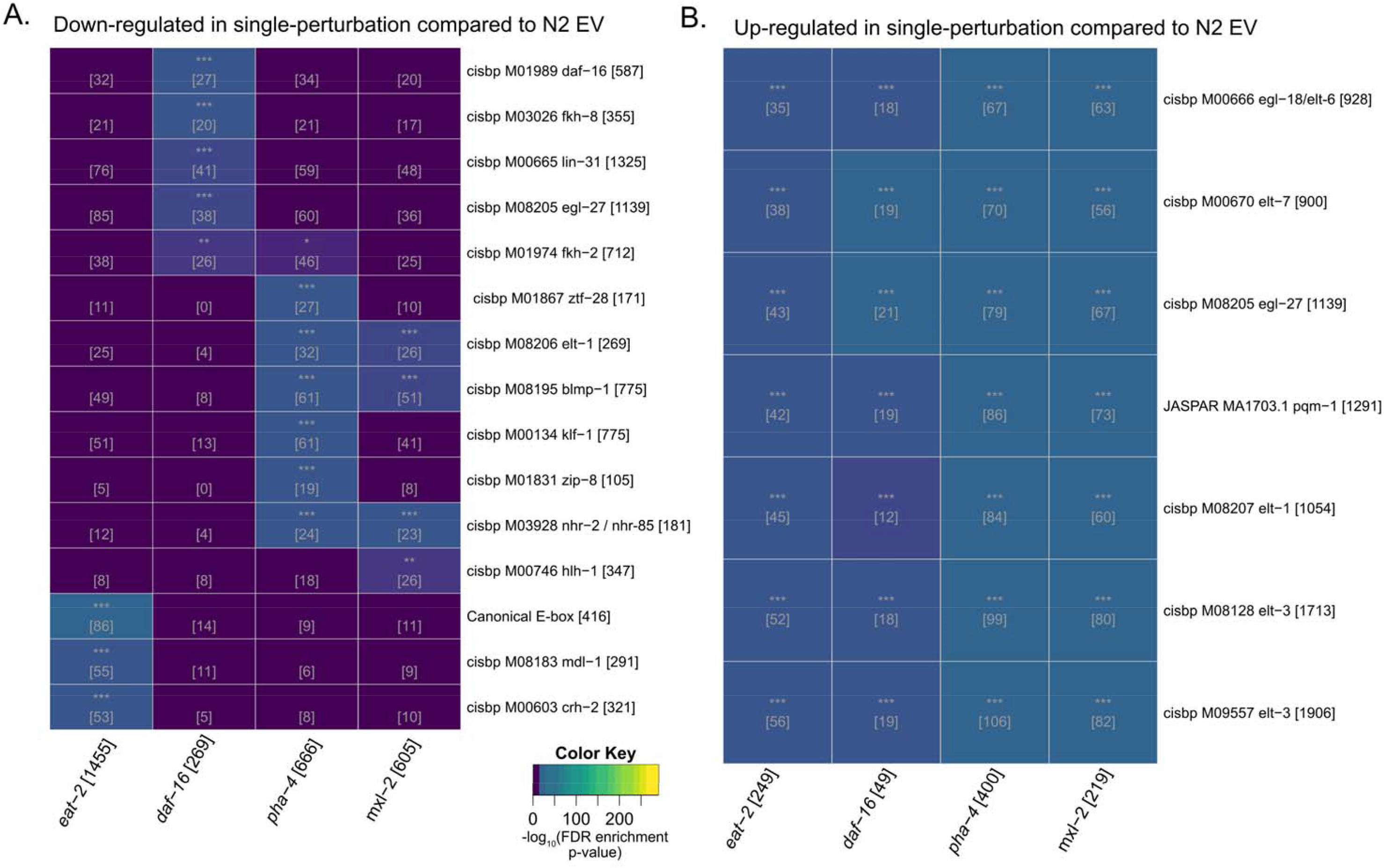
Transcription factor motif enrichment for predicted binding in promoters of genes differentially expressed in *eat-2*, m*xl-2*, *daf-16* (RNAi) or *pha-4* (RNAi) single-perturbation conditions. **(A).** The most enriched motif for genes down-regulated under DR corresponds to canonical CACGTG E-box sites as well as the motif for MDL-1, consistent with the expected binding motif of MML-1::MXL-2. We also see that DAF-16 motifs are most significantly enriched for genes down-regulated with RNAi for *daf-16*. We do not observe enrichment for PHA-4 predicted sites in samples with RNAi for *pha-4*, or in *eat-2* DR conditions. Similarly, we do not see enrichment of E-box sites or motifs of other E-box binding factors enriched in the genes downregulated in *mxl-2* mutants, however MML-1::MXL-2 activity is expected to be low under basal conditions (Johnson et al 2014). **(B).** Surprisingly, the same set of transcription factor motifs represent the most significantly enriched motifs across all single-perturbation conditions for up-regulated genes. These motifs correspond to GATA-binding factors. All represented motif-based binding predictions are after filtering based on matches in homologous genes in two or more other nematode species (see materials and methods for additional detail). The numbers in brackets are the size of the gene set, or the size of the intersection. Stars indicate significant enrichment results. Adjusted p-value significance level: ‘***’ p < 0.001 ‘**’ 0.01 ‘*’ 0.05 ‘.’ 0.1.

## References

Akashi H, Gojobori T (2002) Metabolic efficiency and amino acid composition in the proteomes of Escherichia coli and Bacillus subtilis. Proceedings of the National Academy of Sciences of the United States of America 99:. 10.1073/pnas.062526999

Albert Hubbard EJ (2007) Caenorhabditis elegans germ line: A model for stem cell biology. Dev Dyn 236:3343–3357. 10.1002/dvdy.21335

Allevato M, Bolotin E, Grossman M, et al (2017) Sequence-specific DNA binding by MYC/MAX to low-affinity non-E-box motifs. PLoS ONE 12:1–20. 10.1371/journal.pone.0180147

Andrews S (2010) FastQC: a quality control tool for high throughput sequence data

Antikainen H, Driscoll M, Haspel G (2017) TOR-mediated regulation of metabolism in aging. Aging Cell 16:1219–1233. 10.1111/acel.12689

Apfeld J, O’Connor G, McDonagh T, et al (2005) The AMP-activated protein kinase AAK-2 links energy levels and insulin-like signals to lifespan in C. elegans. Genes and Development 19:411. 10.1101/gad.1255404.3004

Avery L (1993) The genetics of feeding in Caenorhabditis elegans. Genetics 133:. 10.1093/genetics/133.4.897

Bar DZ, Charar C, Dorfman J, et al (2016) Cell size and fat content of dietary-restricted *Caenorhabditis elegans* are regulated by ATX-2, an mTOR repressor. Proceedings of the National Academy of Sciences 113:E4620–E4629. 10.1073/pnas.1512156113

Barbieri M, Bonafè M, Franceschi C, Paolisso G (2003) Insulin/IGF-I-signaling pathway: an evolutionarily conserved mechanism of longevity from yeast to humans. American Journal of Physiology-Endocrinology and Metabolism 285:E1064–E1071. 10.1152/ajpendo.00296.2003

Benian GM, Epstein HF (2011) Caenorhabditis elegans muscle: A genetic and molecular model for protein interactions in the heart. Circulation Research 109:1082–1095. 10.1161/CIRCRESAHA.110.237685

Bennett CF, Vander Wende H, Simko M, et al (2014) Activation of the mitochondrial unfolded protein response does not predict longevity in Caenorhabditis elegans. Nature communications 5:3483. 10.1038/ncomms4483

Berry BJ, Mjelde E, Carreno F, et al (2023) Preservation of mitochondrial membrane potential is necessary for lifespan extension from dietary restriction. GeroScience 1–9

Billin AN, Ayer DE (2006) The Mlx network: Evidence for a parallel max-like transcriptional network that regulates energy metabolism. Current Topics in Microbiology and Immunology 302:255–278. 10.1007/3-540-32952-8_10

Blackwell TK, Sewell AK, Wu Z, Han M (2019) TOR Signaling in *Caenorhabditis elegans* Development, Metabolism, and Aging. Genetics 213:329–360. 10.1534/genetics.119.302504

Blake JA, Dolan M, Drabkin H, et al (2013) Gene ontology annotations and resources. Nucleic Acids Research 41:530–535. 10.1093/nar/gks1050

Blumenthal T, Evans D, Link CD, et al (2002) A global analysis of Caenorhabditis elegans operons. Nature 417:. 10.1038/nature00831

Bolger AM, Lohse M, Usadel B (2014) Trimmomatic: A flexible trimmer for Illumina sequence data. Bioinformatics 30:2114–2120. 10.1093/bioinformatics/btu170

Botts MR, Cohen LB, Probert CS, et al (2016) Microsporidia Intracellular Development Relies on Myc Interaction Network Transcription Factors in the Host. G3&#58; Genes|Genomes|Genetics 6:2707–2716. 10.1534/g3.116.029983

Boyd WA, McBride SJ, Freedman JH (2007) Effects of Genetic Mutations and Chemical Exposures on Caenorhabditis elegans Feeding: Evaluation of a Novel, High-Throughput Screening Assay. PLoS ONE 2:e1259. 10.1371/journal.pone.0001259

Brenner S, Brenner S (1974) THE GENETICS OF CAENORΉABDITIS ELEGANS. Molecular Biology 336:71–94. 10.1002/cbic.200300625

Byerly L, Cassada RC, Russell RL (1976) The life cycle of the nematode Caenorhabditis elegans. Developmental Biology 51:23–33. 10.1016/0012-1606(76)90119-6

Cairo S, Merla G, Urbinati F, et al (2001) WBSCR14, a gene mapping to the Williams – Beuren syndrome deleted region, is a new member of the Mlx transcription factor network. Human Molecular Genetics 10:617–628

Castro-Mondragon JA, Riudavets-Puig R, Rauluseviciute I, et al (2022) JASPAR 2022: The 9th release of the open-access database of transcription factor binding profiles. Nucleic Acids Research 50:D165–D173. 10.1093/nar/gkab1113

Ceballos A, Esse R, Grishok A (2021) The proline-rich domain of MML-1 is biologically important but not required for localization to target promoters. microPublication Biology 1:

Chen D, Zou L (2019) GSE125718-Expression analysis of C. elegans wild-type N2, eat-2, acs-20 and eat-2; acs-20 mutants

Chen J, Caswell-Chen EP (2004) Facultative Vivipary is a Life-History Trait in Caenorhabditis elegans. J Nematol 36:107–113

Choi YJ (2020) Shedding light on the effects of calorie restriction and its mimetics on skin biology. Nutrients 12:. 10.3390/nu12051529

Chu DS, Liu H, Nix P, et al (2006) Sperm chromatin proteomics identifies evolutionarily conserved fertility factors. Nature 443:101–105. 10.1038/nature05050

Conacci-Sorrell M, McFerrin L, Eisenman RN (2014) An overview of MYC and its interactome. Cold Spring Harbor perspectives in medicine 4:1–24. 10.1101/cshperspect.a014357

Cornaro L (1727) Sure and Certain Methods of attaining a Long and Healthful Life.. Daniel Midwinter

Cornaro L (1905) The art of living long. William F. Butler

Cornwell A, Llop JR, Salzman P, et al (2022) The Replica Set Method is a Robust, Accurate, and High-Throughput Approach for Assessing and Comparing Lifespan in C. elegans Experiments. Frontiers in Aging 3:1–21. 10.3389/fragi.2022.861701

Cornwell AB, Llop JR, Salzman P, et al (2018) The Replica Set Method: A High-throughput Approach to Quantitatively Measure Caenorhabditis elegans Lifespan. JoVE: Journal of Visualized Experiments e57819

Cornwell AB, Samuelson AV (2020) Analysis of Lifespan in C. elegans: Low- and High-Throughput Approaches. In: Methods in Molecular Biology

Cox GN, Kusch M, Edgar RS (1981) Cuticle of Caenorhabditis elegans: its isolation and partial characterization. The Journal of cell biology 90:7–17. 10.1083/jcb.90.1.7

Crawford D, Libina N, Kenyon C (2007) Caenorhabditis elegans integrates food and reproductive signals in lifespan determination. Aging Cell 6:715–721. 10.1111/j.1474-9726.2007.00327.x

Cunningham F, Allen JE, Allen J, et al (2022) Ensembl 2022. Nucleic Acids Research 50:D988–D995. 10.1093/nar/gkab1049

Curtis R, O’Connor G, DiStefano PS (2006) Aging networks in Caenorhabditis elegans: AMP-activated protein kinase (aak-2) links multiple aging and metabolism pathways. Aging Cell 5:119–126. 10.1111/j.1474-9726.2006.00205.x

Das R, Melo JA, Thondamal M, et al (2017) The homeodomain-interacting protein kinase HPK-1 preserves protein homeostasis and longevity through master regulatory control of the HSF-1 chaperone network and TORC1-restricted autophagy in Caenorhabditis elegans. PLoS Genetics 1:1–46

Davis P, Zarowiecki M, Arnaboldi V, et al (2022) WormBase in 2022—data, processes, and tools for analyzing *Caenorhabditis elegans*. Genetics 220:iyac003. 10.1093/genetics/iyac003

de Martin X, Sodaei R, Santpere G (2021) Mechanisms of Binding Specificity among bHLH Transcription Factors. IJMS 22:9150. 10.3390/ijms22179150

Denzel MS, Lapierre LR, Mack HID (2018) Emerging topics in C. elegans aging research: Transcriptional regulation, stress response and epigenetics. Mechanisms of Ageing and Development 1–18. 10.1016/j.mad.2018.08.001

Diolaiti D, McFerrin L, Carroll PA, Eisenman RN (2015) Functional interactions among members of the MAX and MLX transcriptional network during oncogenesis. Biochimica et Biophysica Acta - Gene Regulatory Mechanisms 1849:484–500. 10.1016/j.bbagrm.2014.05.016

Dobin A, Davis CA, Schlesinger F, et al (2013) STAR: Ultrafast universal RNA-seq aligner. Bioinformatics 29:15–21. 10.1093/bioinformatics/bts635

Dues DJ, Andrews EK, Schaar CE, et al (2016) Aging causes decreased resistance to multiple stresses and a failure to activate specific stress response pathways. Aging 8:777–795. 10.18632/aging.100939

Dues DJ, Andrews EK, Senchuk MM, Van Raamsdonk JM (2019) Resistance to Stress Can Be Experimentally Dissociated From Longevity. The Journals of Gerontology: Series A 74:1206–1214. 10.1093/gerona/gly213

Duregon E, Pomatto-Watson LCDD, Bernier M, et al (2021) Intermittent fasting: from calories to time restriction. GeroScience 43:1083–1092. 10.1007/s11357-021-00335-z

Edwards C, Canfield J, Copes N, et al (2015) Mechanisms of amino acid-mediated lifespan extension in Caenorhabditis elegans. BMC Genet 16:8. 10.1186/s12863-015-0167-2

Edwards CB, Copes N, Brito AG, et al (2013) Malate and Fumarate Extend Lifespan in Caenorhabditis elegans. PLoS ONE 8:. 10.1371/journal.pone.0058345

Ellis RE, Stanfield GM (2014) The regulation of spermatogenesis and sperm function in nematodes. Seminars in Cell and Developmental Biology 29:. 10.1016/j.semcdb.2014.04.005

Erijman A, Kozlowski L, Sohrabi-Jahromi S, et al (2020) A High-Throughput Screen for Transcription Activation Domains Reveals Their Sequence Features and Permits Prediction by Deep Learning. Molecular Cell 78:890–902.e6. 10.1016/j.molcel.2020.04.020

Essmann CL, Martinez-Martinez D, Pryor R, et al (2020) Mechanical properties measured by atomic force microscopy define health biomarkers in ageing C. elegans. Nature Communications 11:. 10.1038/s41467-020-14785-0

Ewald CY, Landis JN, Abate JP, et al (2015) Dauer-independent insulin/IGF-1-signalling implicates collagen remodelling in longevity. Nature 519:97–101. 10.1038/nature14021

Fontana L, Partridge L, Longo VD (2010) Extending healthy life span - From yeast to humans. Science 328:321–326. 10.1126/science.1172539

Fraga D, Aryal M, Hall JE, et al (2015) Characterization of the arginine kinase isoforms in Caenorhabditis elegans. Comparative Biochemistry and Physiology Part B: Biochemistry and Molecular Biology 187:85–101. 10.1016/j.cbpb.2015.05.002

Fuxman Bass JI, Pons C, Kozlowski L, et al (2016) A gene-centered C. elegans protein–DNA interaction network provides a framework for functional predictions. Molecular Systems Biology 12:1–19. 10.15252/msb.20167131

Gao AW, Smith RL, Weeghel MV, et al (2018) Identification of key pathways and metabolic fingerprints of longevity in C . elegans. Experimental Gerontology 113:128–140. 10.1101/222554

Gao AW, uit de Bos J, Sterken MG, et al (2017) Forward and reverse genetics approaches to uncover metabolic aging pathways in Caenorhabditis elegans. Biochimica et Biophysica Acta - Molecular Basis of Disease 0–1. 10.1016/j.bbadis.2017.09.006

Gelino S, Chang JT, Kumsta C, et al (2016) Intestinal Autophagy Improves Healthspan and Longevity in C. elegans during Dietary Restriction. PLoS Genetics 12:1–24. 10.1371/journal.pgen.1006135

Gerstein MB, Lu ZJ, Van Nostrand EL, et al (2010) Integrative analysis of the Caenorhabditis elegans genome by the modENCODE project. Science 330:1775–1787. 10.1126/science.1196914

Gilbert W, Guthrie C (2004) The Glc7p Nuclear Phosphatase Promotes mRNA Export by Facilitating Association of Mex67p with mRNA. Molecular Cell 13:201–212. 10.1016/S1097-2765(04)00030-9

Gillespie M, Jassal B, Stephan R, et al (2022) The reactome pathway knowledgebase 2022. 50:687–692

Glenwinkel L, Taylor SR, Langebeck-Jensen K, et al (2021) In silico analysis of the transcriptional regulatory logic of neuronal identity specification throughout the c. Elegans nervous system. eLife 10:1–29. 10.7554/eLife.64906

Gouvea DY, Aprison EZ, Ruvinsky I (2015) Experience Modulates the Reproductive Response to Heat Stress in C . elegans via Multiple Physiological Processes. 1–27. 10.1371/journal.pone.0145925

Grant CE, Bailey TL, Noble WS (2011) FIMO: Scanning for occurrences of a given motif. Bioinformatics 27:1017–1018. 10.1093/bioinformatics/btr064

Green CL, Lamming DW, Fontana L (2022) Molecular mechanisms of dietary restriction promoting health and longevity. Nature Reviews Molecular Cell Biology 23:56–73. 10.1038/s41580-021-00411-4

Greer EL, Brunet A (2009) Different dietary restriction regimens extend lifespan by both independent and overlapping genetic pathways in C. elegans. Aging Cell 8:113–127. 10.1111/j.1474-9726.2009.00459.x

Greer EL, Dowlatshahi D, Banko MR, et al (2007) An AMPK-FOXO Pathway Mediates Longevity Induced by a Novel Method of Dietary Restriction in C. elegans. Current Biology 17:1646–1656. 10.1016/j.cub.2007.08.047

Grove CA, De Masi F, Barrasa MI, et al (2009) A Multiparameter Network Reveals Extensive Divergence between C. elegans bHLH Transcription Factors. Cell 138:314–327. 10.1016/j.cell.2009.04.058

Hamilton B, Dong Y, Shindo M, et al (2005) A systematic RNAi screen for longevity genes in C. elegans. Genes and Development 19:1544–1555. 10.1101/gad.1308205

Hansen M, Chandra A, Mitic LL, et al (2008) A role for autophagy in the extension of lifespan by dietary restriction in C. elegans. PLoS Genetics 4:. 10.1371/journal.pgen.0040024

Hansen M, Hsu AL, Dillin A, Kenyon C (2005) New genes tied to endocrine, metabolic, and dietary regulation of lifespan from a Caenorhabditis elegans genomic RNAi screen. PLoS Genetics 1:0119–0128. 10.1371/journal.pgen.0010017

Hansen M, Taubert S, Crawford D, et al (2007) Lifespan extension by conditions that inhibit translation in *Caenorhabditis elegans*. Aging Cell 6:95–110. 10.1111/j.1474-9726.2006.00267.x

Harris JE, Govindan JA, Yamamoto I, et al (2006) Major sperm protein signaling promotes oocyte microtubule reorganization prior to fertilization in Caenorhabditis elegans. Developmental Biology 299:105–121. 10.1016/j.ydbio.2006.07.013

Harris TW, Antoshechkin I, Bieri T, et al (2009) Wormbase: A comprehensive resource for nematode research. Nucleic Acids Research 38:463–467. 10.1093/nar/gkp952

Heestand BN, Shen Y, Liu W, et al (2013) Dietary restriction induced longevity is mediated by nuclear receptor NHR-62 in Caenorhabditis elegans. PLoS genetics 9:e1003651. 10.1371/journal.pgen.1003651

Henderson ST, Bonafè M, Johnson TE (2006) daf-16 protects the nematode Caenorhabditis elegans during food deprivation. Journals of Gerontology - Series A Biological Sciences and Medical Sciences 61:444–460. 10.1093/gerona/61.5.444

Hofmann JW, Zhao X, De Cecco M, et al (2015) Reduced Expression of MYC Increases Longevity and Enhances Healthspan. Cell 160:477–488. 10.1016/j.cell.2014.12.016

Horowitz BB, Nanda S, Walhout AJM (2023) A transcriptional cofactor regulatory network for the C. elegans intestine. G3 (Bethesda) 13:jkad096. 10.1093/g3journal/jkad096

Houthoofd K, Braeckman BP, Lenaerts I, et al (2002) No reduction of metabolic rate in food restricted Caenorhabditis elegans. 37:1357–1367

Hume MA, Barrera LA, Gisselbrecht SS, Bulyk ML (2015) UniPROBE, update 2015: New tools and content for the online database of protein-binding microarray data on protein-DNA interactions. Nucleic Acids Research 43:D117–D122. 10.1093/nar/gku1045

Il’yasova D, Fontana L, Bhapkar M, et al (2018) Effects of 2 years of caloric restriction on oxidative status assessed by urinary F2-isoprostanes: The CALERIE 2 randomized clinical trial. Aging Cell 17:1–9. 10.1111/acel.12719

Izaurralde E (2004) Directing mRNA export. Nat Struct Mol Biol 11:210–212. 10.1038/nsmb0304-210

Jedrusik-Bode M (2014) *C. elegans* sirtuin SIR-2.4 and its mammalian homolog SIRT6 in stress response. Worm 3:e29102. 10.4161/worm.29102

Jedrusik-Bode M, Studencka M, Smolka C, et al (2013) The sirtuin SIRT6 regulates stress granules formation in *C. elegans* and in mammals. Journal of Cell Science jcs.130708. 10.1242/jcs.130708

Jia K, Chen D, Riddle DL (2004) The TOR pathway interacts with the insulin signaling pathway to regulate C. elegans larval development, metabolism and life span. Development 131:3897–3906. 10.1242/dev.01255

Johnson DW, Llop JR, Farrell SF, et al (2014) The Caenorhabditis elegans Myc-Mondo/Mad Complexes Integrate Diverse Longevity Signals. PLoS Genetics 10:e1004278. 10.1371/journal.pgen.1004278

Johnson T (1990) Increased life-span of age-1 mutants in Caenorhabditis elegans and lower Gompertz rate of aging. Science 249:908–912. 10.1126/science.2392681

Johnson TE, Wood WB (1982) Genetic analysis of life-span in Caenorhabditis elegans. Proceedings of the National Academy of Sciences of the United States of America 79:6603–7. 10.1073/pnas.79.21.6603

Johnstone IL (2000) Cuticle collagen genes expression in Caenorhabditis elegans Collagen is a structural protein used in the generation of a wide variety of animal extracellular matrices . The. Trends in Genetics 16:21–27

Kaeberlein TL, Smith ED, Tsuchiya M, et al (2006) Lifespan extension in Caenorhabditis elegans by complete removal of food. Aging Cell 5:487–494. 10.1111/j.1474-9726.2006.00238.x

Kaletsky R, Yao V, Williams A, et al (2018) Transcriptome analysis of adult Caenorhabditis elegans cells reveals tissue-specific gene and isoform expression. PLoS Genetics 14:. 10.1371/journal.pgen.1007559

Kamath RS, Martinez-Campos M, Zipperlen P, et al (2000) Effectiveness of specific RNA-mediated interference through ingested double-stranded RNA in Caenorhabditis elegans. Genome Biology 2:research0002.1–10. 10.1186/gb-2000-2-1-research0002

Kanehisa M, Goto S, Furumichi M, et al (2010) KEGG for representation and analysis of molecular networks involving diseases and drugs. Nucleic acids research 38:D355–60. 10.1093/nar/gkp896

Kapahi P, Chen D, Rogers AN, et al (2010) With TOR, Less Is More: A Key Role for the Conserved Nutrient-Sensing TOR Pathway in Aging. Cell Metabolism 11:453–465. 10.1016/j.cmet.2010.05.001

Kapahi P, Kaeberlein M, Hansen M (2017) Dietary restriction and lifespan: Lessons from invertebrate models. Ageing Research Reviews 39:3–14. 10.1016/j.arr.2016.12.005

Kennedy BK, Steffen KK, Kaeberlein M (2007) Ruminations on dietary restriction and aging. Cellular and Molecular Life Sciences 64:1323–1328. 10.1007/s00018-007-6470-y

Kenyon C, Chang J, Gensch E, et al (1993) A C. elegans mutant that lives twice as long as wild type. Nature 366:461–464. 10.1038/366461a0

Kim W, Underwood RS, Greenwald I, Shaye DD (2018) Ortholist 2: A new comparative genomic analysis of human and caenorhabditis elegans genes. Genetics 210:445–461. 10.1534/genetics.118.301307

Kniazeva M, Crawford QT, Seiber M, et al (2004) Monomethyl branched-chain fatty acids play an essential role in Caenorhabditis elegans development. PLoS Biology 2:. 10.1371/journal.pbio.0020257

Korta DZ, Tuck S, Hubbard EJA (2012) S6K links cell fate, cell cycle and nutrient response in C. elegans germline stem/progenitor cells. Development 139:859–870. 10.1242/dev.074047

Kraus WE, Bhapkar M, Huffman KM, et al (2019) 2 years of calorie restriction and cardiometabolic risk (CALERIE): exploratory outcomes of a multicentre, phase 2, randomised controlled trial. The Lancet Diabetes and Endocrinology 7:673–683. 10.1016/S2213-8587(19)30151-2

Krittika S, Yadav P (2019) An overview of two decades of diet restriction studies using Drosophila. Biogerontology 20:723–740. 10.1007/s10522-019-09827-0

Kumar S, Egan BM, Kocsisova Z, et al (2019) Lifespan Extension in C. elegans Caused by Bacterial Colonization of the Intestine and Subsequent Activation of an Innate Immune Response. Developmental Cell 49:100–117.e6. 10.1016/j.devcel.2019.03.010

Labbadia J, Morimoto RI (2015) Repression of the Heat Shock Response Is a Programmed Event at the Onset of Reproduction. Molecular Cell 59:639–650. 10.1016/j.molcel.2015.06.027

Lahoz EG, Xu L, Schreiber-Agus N, DePinho RA (1994) Suppression of Myc, but not E1a, transformation activity by Max-associated proteins, Mad and Mxi1. Proc Natl Acad Sci USA 91:5503–5507. 10.1073/pnas.91.12.5503

Lakowski B, Hekimi S (1998) The genetics of caloric restriction in Caenorhabditis elegans. Proc Natl Acad Sci USA 95:13091–13096

Lapierre LR, De Magalhaes Filho CD, McQuary PR, et al (2013) The TFEB orthologue HLH-30 regulates autophagy and modulates longevity in Caenorhabditis elegans. Nature Communications 4:1–8. 10.1038/ncomms3267

Lapierre LR, Kumsta C, Sandri M, et al (2015) Transcriptional and epigenetic regulation of autophagy in aging. Autophagy 11:867–880. 10.1080/15548627.2015.1034410

Laplante M, Sabatini DM (2012) Review mTOR Signaling in Growth Control and Disease. CELL 149:274–293. 10.1016/j.cell.2012.03.017

Lazaro-Pena MI, Cornwell AB, Diaz-Balzac CA, et al (2023) Homeodomain-interacting protein kinase maintains neuronal homeostasis during normal Caenorhabditis elegans aging and systemically regulates longevity from serotonergic and GABAergic neurons. Elife 12:e85792

Lazaro-Pena MI, Ward ZC, Yang S, et al (2022) HSF-1: Guardian of the Proteome Through Integration of Longevity Signals to the Proteostatic Network. Front Aging 3:861686. 10.3389/fragi.2022.861686

Lee S, Kenyon C (2009) Article Regulation of the Longevity Response to Temperature by Thermosensory Neurons in Caenorhabditis elegans. Current Biology 19:715–722. 10.1016/j.cub.2009.03.041

Lee SJ, Hwang AB, Kenyon C (2010) Inhibition of respiration extends C. elegans life span via reactive oxygen species that increase HIF-1 activity. Current Biology 20:2131–2136. 10.1016/j.cub.2010.10.057

Li B, Dewey CN (2011) RSEM: accurate transcript quantification from RNA-Seq data with or without a reference genome. BMC Bioinformatics 12:323. 10.1186/1471-2105-12-323

Liao Y, Smyth GK, Shi W (2014) FeatureCounts: An efficient general purpose program for assigning sequence reads to genomic features. Bioinformatics 30:923–930. 10.1093/bioinformatics/btt656

Lim YP, Go MK, Raida M, et al (2018) Synthetic Enzymology and the Fountain of Youth: Repurposing Biology for Longevity. ACS Omega 3:. 10.1021/acsomega.8b01620

Lin K, Dorman JB, Rodan A, Kenyon C (1997) daf-16: An HNF-3/forkhead family member that can function to double the life-span of Caenorhabditis elegans. Science 278:1319–1322. 10.1126/science.278.5341.1319

Lin MG, Hurley JH (2016) Structure and function of the ULK1 complex in autophagy. Current Opinion in Cell Biology 39:61–68. 10.1016/j.ceb.2016.02.010

Love MI, Huber W, Anders S (2014) Moderated estimation of fold change and dispersion for RNA-seq data with DESeq2. Genome biology 15:550. 10.1186/s13059-014-0550-8

Luo S, Shaw WM, Ashraf J, Murphy CT (2009) TGF-ß Sma/Mab signaling mutations uncouple reproductive aging from somatic aging. PLoS Genetics 5:. 10.1371/journal.pgen.1000789

Mair W, Dillin A (2008) Aging and Survival: The Genetics of Life Span Extension by Dietary Restriction. Annual Review of Biochemistry 77:727–754. 10.1146/annurev.biochem.77.061206.171059

Mair W, Panowski SH, Shaw RJ, Dillin A (2009) Optimizing dietary restriction for genetic epistasis analysis and gene discovery in C. elegans. PLoS ONE 4:. 10.1371/journal.pone.0004535

Matai L, Sarkar GC, Chamoli M, et al (2019) Dietary restriction improves proteostasis and increases life span through endoplasmic reticulum hormesis. Proceedings of the National Academy of Sciences of the United States of America 116:17383–17392. 10.1073/pnas.1900055116

Mattison JA, Colman RJ, Beasley TM, et al (2017) Caloric restriction improves health and survival of rhesus monkeys. Nature Communications 8:14063. 10.1038/ncomms14063

McFerrin LG, Atchley WR (2011) Evolution of the max and MlX networks in animals. Genome Biology and Evolution 3:915–937. 10.1093/gbe/evr082

Mcferrin LG, Atchley WR (2012) A Novel N-Terminal Domain May Dictate the Glucose Response of Mondo Proteins. 7:1–16. 10.1371/journal.pone.0034803

Mckay JP, Raizen DM, Gottschalk A, et al (2004) eat-2 and eat-18 Are Required for Nicotinic Neurotransmission in the Caenorhabditis elegans Pharynx. Genetics 166:161–169

McQuary PR, Liao C-Y, Chang JT, et al (2016) C. elegans S6K Mutants Require a Creatine-Kinase-like Effector for Lifespan Extension. Cell Reports 14:2059–2067. 10.1016/j.celrep.2016.02.012

Meléndez A, Levine B (2009) Autophagy in C. elegans. WormBook: the online review of C elegans biology 1–26

Mesbahi H, Pho KB, Tench AJ, et al (2020) Cuticle collagen expression is regulated in response to environmental stimuli by the GATA transcription factor ELT-3 in caenorhabditis elegans. Genetics 215:483–495. 10.1534/genetics.120.303125

Miller HA, Dean ES, Pletcher SD, Leiser SF (2020) Cell non-autonomous regulation of health and longevity. eLife 9:e62659. 10.7554/eLife.62659

Mouchiroud L, Molin L, Mergoud-dit-lamarche A (2015) Metabolomics Analysis Uncovers That Dietary Restriction Buffers Metabolic Changes Associated with Aging in Caenorhabditis elegans. Journal of Proteome Research 13:2910–2919

Mukhopadhyay A, Oh SW, Tissenbaum HA (2006) Worming pathways to and from DAF-16/FOXO. Experimental Gerontology 41:928–934. 10.1016/j.exger.2006.05.020

Murphy CT, McCarroll S a, Bargmann CI, et al (2003) Genes that act downstream of DAF-16 to influence the lifespan of Caenorhabditis elegans. Nature 424:277–283. 10.1038/nature01789

Nakagawa S, Lagisz M, Hector KL, Spencer HG (2012) Comparative and meta-analytic insights into life extension via dietary restriction. Aging Cell 11:401–409. 10.1111/j.1474-9726.2012.00798.x

Nakamura S, Karalay Ö, Jäger PS, et al (2016) Mondo complexes regulate TFEB via TOR inhibition to promote longevity in response to gonadal signals. Nature communications 7:10944. 10.1038/ncomms10944

Ogg S, Paradis S, Gottlieb S, et al (1997) The fork head transcription factor DAF-16 transduces insulin-like metabolic and longevity signals in C. elegans. Nature 389:994–999. 10.1038/40194

O’Rourke EJ, Ruvkun G (2013) MXL-3 and HLH-30 transcriptionally link lipolysis and autophagy to nutrient availability. Nature cell biology 15:668–76. 10.1038/ncb2741

Pandit A, Jain V, Kumar N, Mukhopadhyay A (2014) PHA-4/FOXA-regulated microRNA feed forward loops during Caenorhabditis elegans dietary restriction. Aging 6:835–855. https://doi.org/100697 [pii]

Pang S, Lynn DA, Lo JY, et al (2014) SKN-1 and Nrf2 couples proline catabolism with lipid metabolism during nutrient deprivation. Nat Commun 5:5048. 10.1038/ncomms6048

Panowski SH, Wolff S, Aguilaniu H, et al (2007) PHA-4/Foxa mediates diet-restriction-induced longevity of C. elegans. Nature 447:550–555. 10.1038/nature05837

Paraskevopoulou MD, Georgakilas G, Kostoulas N, et al (2013) DIANA-microT web server v5.0: service integration into miRNA functional analysis workflows. Nucleic acids research 41:169–173. 10.1093/nar/gkt393

Pickett CL, Breen KT, Ayer DE (2007) A C. elegans Myc-like network cooperates with semaphorin and Wnt signaling pathways to control cell migration. Developmental Biology 310:226–239. 10.1016/j.ydbio.2007.07.034

R Core Team R: A Language and Environment for Statistical Computing

Rahimi M, Sohrabi S, Murphy CT (2022) Novel elasticity measurements reveal C. elegans cuticle stiffens with age and in a long-lived mutant. Biophysical Journal 121:515–524. 10.1016/j.bpj.2022.01.013

Reinke V, Smith HE, Nance J, et al (2000) A global profile of germline gene expression in C. elegans. Molecular Cell 6:. 10.1016/S1097-2765(00)00059-9

Rodríguez-Palero MJ, López-Díaz A, Marsac R, et al (2018) An automated method for the analysis of food intake behaviour in Caenorhabditis elegans. Scientific Reports 8:1–10. 10.1038/s41598-018-21964-z

Saito TL, Hashimoto SI, Gu SG, et al (2013) The transcription start site landscape of C. elegans. Genome Research 23:1348–1361. 10.1101/gr.151571.112

Salminen A, Kaarniranta K (2012) AMP-activated protein kinase (AMPK) controls the aging process via an integrated signaling network. Ageing Research Reviews 11:230–241. 10.1016/j.arr.2011.12.005

Samuelson AV, Carr CE, Ruvkun G (2007) Gene activities that mediate increased life span of C. elegans insulin-like signaling mutants. Genes & Development 21:2976–94. 10.1101/gad.1588907

Sanborn AL, Yeh BT, Feigerle JT, et al (2021) Simple biochemical features underlie transcriptional activation domain diversity and dynamic, fuzzy binding to mediator. eLife 10:1–42. 10.7554/ELIFE.68068

Sandhu A, Badal D, Sheokand R, et al (2021) Specific collagens maintain the cuticle permeability barrier in Caenorhabditis elegans. Genetics 217:. 10.1093/GENETICS/IYAA047

Scharf A, Pohl F, Egan BM, et al (2021) Reproductive Aging in Caenorhabditis elegans: From Molecules to Ecology. Front Cell Dev Biol 9:718522. 10.3389/fcell.2021.718522

Schreiber-Agus N, Chin L, Chen K, et al (1994) Evolutionary relationships and functional conservation among vertebrate Max-associated proteins: the zebra fish homolog of Mxi1. Oncogene 9:3167—3177

Seo K, Choi E, Lee D, et al (2013) Heat shock factor 1 mediates the longevity conferred by inhibition of TOR and insulin/IGF-1 signaling pathways in C. elegans. Aging Cell 12:1073–1081. 10.1111/acel.12140

Sharples AP, Hughes DC, Deane CS, et al (2015) Longevity and skeletal muscle mass: the role of IGF signalling, the sirtuins, dietary restriction and protein intake. Aging Cell 14:511–523. 10.1111/acel.12342

Shaw WM, Luo S, Landis J, et al (2007) The C. elegans TGF-B Dauer Pathway Regulates Longevity via Insulin Signaling. Current Biology 17:1635–1645. 10.1016/j.cub.2007.08.058

Sheaffer KL, Updike DL, Mango SE (2008) The Target of Rapamycin Pathway Antagonizes pha-4/FoxA to Control Development and Aging. Current Biology 18:1355–1364. 10.1016/j.cub.2008.07.097

Shih HM, Liu Z, Towle HC (1995) Two CACGTG motifs with proper spacing dictate the carbohydrate regulation of hepatic gene transcription. Journal of Biological Chemistry 270:21991–21997. 10.1074/jbc.270.37.21991

Shimabukuro K, Roberts TM (2013) Major Sperm Protein and Sperm Locomotion. Elsevier Inc.

Shioda T, Takahashi I, Ikenaka K, et al (2023) Neuronal MML-1/MXL-2 regulates systemic aging via glutamate transporter and cell nonautonomous autophagic and peroxidase activity. Proc Natl Acad Sci USA 120:e2221553120. 10.1073/pnas.2221553120

Shpigel N, Shemesh N, Kishner M, Ben-Zvi A (2019) Dietary restriction and gonadal signaling differentially regulate post-development quality control functions in Caenorhabditis elegans. Aging Cell 18:e12891. 10.1111/acel.12891

Singson A (2001) Every Sperm Is Sacred: Fertilization in Caenorhabditis elegans. Developmental Biology 230:101–109. 10.1006/dbio.2000.0118

Sloan EJ, Ayer DE (2010) Myc, Mondo, and Metabolism. Genes & Cancer 1:587–596. 10.1177/1947601910377489

Sohal RS, Weindruch R (1996) Oxidative stress, caloric restriction, and aging. Science 273:59–63. 10.1126/science.273.5271.59

Son HG, Altintas O, Kim EJE, et al (2019) Age-dependent changes and biomarkers of aging in Caenorhabditis elegans. Aging Cell 18:1–11. 10.1111/acel.12853

Soo SK, Traa A, Rudich ZD, et al (2023) Genetic basis of enhanced stress resistance in long-lived mutants highlights key role of innate immunity in determining longevity. Aging Cell 22:. 10.1111/acel.13740

Soukas AA, Kane EA, Carr CE, et al (2009) Rictor / TORC2 regulates fat metabolism, feeding, growth, and life span in Caenorhabditis elegans. Genes and Development 23:496–511. 10.1101/gad.1775409.2004

Soultoukis GA, Partridge L (2016) Dietary Protein, Metabolism, and Aging. Annu Rev Biochem 85:5–34. 10.1146/annurev-biochem-060815-014422

Souza Matos M, Platt B, Delibegovic M (2021) Effects of dietary restriction on metabolic and cognitive health. Proceedings of the Nutrition Society 80:126–138. 10.1017/S0029665120007910

Steinkraus KA, Smith ED, Davis C, et al (2008) Dietary restriction suppresses proteotoxicity and enhances longevity by an hsf-1-dependent mechanism in Caenorhabditis elegans. Aging Cell 7:394–404. 10.1111/j.1474-9726.2008.00385.x

Stekovic S, Hofer SJ, Tripolt N, et al (2019) Alternate Day Fasting Improves Physiological and Molecular Markers of Aging in Healthy, Non-obese Humans. Cell Metabolism 1–15. 10.1016/j.cmet.2019.07.016

Stroustrup N, Ulmschneider BE, Nash ZM, et al (2014) The C. elegans Lifespan Machine. 10:665– 670. 10.1038/nmeth.2475.The

Sulston JE, Horvitz HR (1977) Post-embryonic cell lineages of the nematode, Caenorhabditis elegans. Developmental Biology 56:. 10.1016/0012-1606(77)90158-0

Tabrez SS, Sharma RD, Jain V, et al (2017) Differential alternative splicing coupled to nonsense-mediated decay of mRNA ensures dietary restriction-induced longevity. Nature Communications 8:306. 10.1038/s41467-017-00370-5

Tacutu R, Thornton D, Johnson E, et al (2018) Human Ageing Genomic Resources: New and updated databases. Nucleic Acids Research 46:D1083–D1090. 10.1093/nar/gk×1042

Taguchi A, White MF (2008) Insulin-like signaling, nutrient homeostasis, and life span. Annual Review of Physiology 70:191–212. 10.1146/annurev.physiol.70.113006.100533

Templeman NM, Murphy CT (2018) Regulation of reproduction and longevity by nutrient-sensing pathways. Journal of Cell Biology 217:93–106. 10.1083/jcb.201707168

Tepper RG, Ashraf J, Kaletsky R, et al (2013) PQM-1 Complements DAF-16 as a Key Transcriptional Regulator of DAF-2-Mediated Development and Longevity. Cell 154:676–690. 10.1016/j.cell.2013.07.006

Timmons L, Fire A (1998) Specific interference by ingested dsRNA [10]. Nature 395:854. 10.1038/27579

Tóth ML, Sigmond T, Borsos É, et al (2008) Longevity pathways converge on autophagy genes to regulate life span in Caenorhabditis elegans. Autophagy 8627:. 10.4161/auto.5618

Tullet JMA, Hertweck M, An JH, et al (2008) Direct Inhibition of the Longevity-Promoting Factor SKN-1 by Insulin-like Signaling in C. elegans. Cell 132:1025–1038. 10.1016/j.cell.2008.01.030

Verdin E (2015) NAD ^+^ in aging, metabolism, and neurodegeneration. Science 350:1208–1213. 10.1126/science.aac4854

Vieira AFC, Xatse MA, Tifeki H, et al (2022) Monomethyl branched-chain fatty acids are critical for Caenorhabitis elegans survival in elevated glucose conditions. Journal of Biological Chemistry 298:101444. 10.1016/j.jbc.2021.101444

von Frieling J, Roeder T (2020) Factors that affect the translation of dietary restriction into a longer life. IUBMB Life 72:814–824. 10.1002/iub.2224

Vos MJ, Carra S, Kanon B, et al (2016) Specific protein homeostatic functions of small heat-shock proteins increase lifespan. Aging Cell 15:217–226. 10.1111/acel.12422

Weir HJ, Yao P, Huynh FK, et al (2017) Dietary Restriction and AMPK Increase Lifespan via Mitochondrial Network and Peroxisome Remodeling. Cell Metabolism 26:884–896.e5. 10.1016/j.cmet.2017.09.024

Willcox BJ, Donlon TA, He Q, et al (2008) FOXO3A genotype is strongly associated with human longevity. Proceedings of the National Academy of Sciences of the United States of America 105:13987– 13992. 10.1073/pnas.0801030105

Yang W, Dierking K, Schulenburg H (2016) WormExp: a web-based application for a Caenorhabditis elegans-specific gene expression enrichment analysis. Bioinformatics (Oxford, England) 32:943–5. 10.1093/bioinformatics/btv667

Young MD, Wakefield MJ, Smyth GK, Oshlack A (2010) Gene ontology analysis for RNA-seq: accounting for selection bias. Genome biology 11:R14. 10.1186/gb-2010-11-2-r14

Yuan J, Tirabassi RS, Bush a B, Cole MD (1998) The C. elegans MDL-1 and MXL-1 proteins can functionally substitute for vertebrate MAD and MAX. Oncogene 17:1109–1118. 10.1038/sj.onc.1202036

Yuan Y, Kadiyala CS, Ching T, et al (2012) Enhanced Energy Metabolism Contributes to the Extended Life Span of Calorie-restricted Caenorhabditis elegans. The Journal of biological chemistry 287:31414–31426. 10.1074/jbc.M112.377275

Zaret KS, Carroll JS (2011) Pioneer transcription factors: establishing competence for gene expression. Genes Dev 25:2227–2241. 10.1101/gad.176826.111

Zaret KS, Mango SE (2016) Pioneer transcription factors, chromatin dynamics, and cell fate control. Current opinion in genetics & development 37:76–81. 10.1016/j.gde.2015.12.003

Zarse K, Schmeisser S, Groth M, et al (2012) Impaired Insulin/IGF1 Signaling Extends Life Span by Promoting Mitochondrial L-Proline Catabolism to Induce a Transient ROS Signal. Cell Metabolism 15:451–465. 10.1016/j.cmet.2012.02.013

Zečić A, Dhondt I, Braeckman BP (2019) The nutritional requirements of Caenorhabditis elegans. Genes and Nutrition 14:. 10.1186/s12263-019-0637-7

Zhang L, Ward JD, Cheng Z, Dernburg AF (2015) The auxin-inducible degradation (AID) system enables versatile conditional protein depletion in C. elegans. Development (Cambridge) 142:4374–4384. 10.1242/dev.129635

Zhang Y, Shao Z, Zhai Z, et al (2009) The HIF-1 Hypoxia-Inducible Factor Modulates Lifespan in C. elegans. PLoS ONE 4:e6348. 10.1371/journal.pone.0006348

Zhao Y, Wang H, Poole RJ, Gems D (2019) A fln-2 mutation affects lethal pathology and lifespan in C. elegans. Nature Communications 10:1–10. 10.1038/s41467-019-13062-z

Zhong M, Niu W, Lu ZJ, et al (2010) Genome-Wide Identification of Binding Sites Defines Distinct Functions for Caenorhabditis elegans PHA-4 / FOXA in Development and Environmental Response. 6:. 10.1371/journal.pgen.1000848

